# Mesenchymal stromal cell remodeling of a gelatin hydrogel microenvironment defines an artificial hematopoietic stem cell niche

**DOI:** 10.1101/289553

**Authors:** Aidan E. Gilchrist, Sunho Lee, Yuhang Hu, Brendan A.C. Harley

## Abstract

Hematopoietic stem cells (HSCs) reside in the bone marrow within discrete niches defined by a complex milieu of external signals including biophysical cues, bound and diffusible biomolecules, and heterotypic cell-cell interactions. Recent studies have shown the importance of autocrine-mediated feedback of cell-secreted signals and the interplay between matrix architecture and biochemical diffusion on hematopoietic stem cell activity. Autocrine and paracrine signaling from HSCs and niche-associated mesenchymal stromal cells (MSCs) have both been suggested to support HSC maintenance *in vivo* and *in vitro*. Here we report the development of a library of methacrylamide-functionalized gelatin (GelMA) hydrogels to explore the balance between autocrine feedback and paracrine signals from co-encapsulated murine bone marrow MSCs on murine HSCs. The use of a degradable GelMA hydrogel enables the possibility for significant MSC-mediated remodeling, yielding dynamic shifts in the matrix environment surrounding HSCs. We identify a combination of an initially low-diffusivity hydrogel and a 1:1 HSPC:MSC seeding ratio as conducive to enhanced HSC population maintenance and quiescence. Further, gene expression and serial mechanical testing data suggests that MSC-mediated matrix remodeling is significant for the long-term HSC culture, reducing HSC autocrine feedback and potentially enhancing MSC-mediated signaling over 7-day culture *in vitro*. This work demonstrates the design of an HSC culture system that couples initial hydrogel properties, MSC co-culture, and concepts of dynamic reciprocity mediated by MSC remodeling to achieve enhanced HSC maintenance.

**One Sentence Summary:** Coupling effects of hydrogel biotransport, heterotypic cell culture, and matrix remodeling enhances hematopoietic stem cell culture and quiescence.

## Introduction

Hematopoietic stem cells (HSCs) possess the capacity for self-renewal and can repopulate the body’s entire complement of blood and immune cells through a highly regulated differentiation hierarchy (*1-3*). Following development, HSCs lodge primarily in the bone marrow in unique tissue microenvironments termed *niches*, which provide a milieu of extrinsic signals in the form of biophysical cues, biomolecular gradients, and co-inhabiting niche cells that together promote quiescence, self-renewal, and lineage specification (*4-9*). Any disruptions to the niche can lead to a loss regulation and abnormal or unsuccessful hemostasis in the bone marrow. The resulting drop in HSC efficacy from niche disruption has hindered clinical use of HSC transplants (HSCTs) for the treatment of blood and immune disorders (*10, 11*). There is an acute clinical need for *in vitro* approaches to improve HSC expansion or preconditioning to facilitate improved homing and engraftment of transplanted donor HSCs to recipient bone marrow niches. Such investigations require biomaterial culture systems that can provide the correct sequence of signals to enhance HSC expansion *in vitro*. While exact replication of the native niche is prohibitively complex due to its transient nature, heterogeneity, and coupling of external factors, insight from the native niche can inform the development of a synthetic culture platform that mediates HSC response (*12*).

Knockout (depletion) and proximity principal (localization) models, both *in vivo* and *in vitro*, have been used to elucidate a range of triggers present in the microenvironment that maintain HSC hematopoietic activity (*13*). *In vivo*, HSCs are presented with complex patterns of ligands such as laminins and fibronectin (*14, 15*) and an effective Young’s modulus ranging from 0.25 to 24.7 kPa (*16*). Matrix biophysical cues within the bone marrow such as elasticity and substrate stiffness can also significantly influence HSC expansion and migration (*17-20*). More recently, tissue engineering approaches have begun to identify critical biomaterial design parameters inspired by native niche signals for *ex vivo* HSC culture. Notably, elastic substrates functionalized with fibronectin have been shown to induce *ex vivo* expansion of HSC progenitor populations (*21*). However, in addition to direct effects of the niche microenvironment on HSC activity, it is increasingly important to examine indirect effects as well. For example, the niche microenvironment can modulate the behavior of co-inhabiting niche cells, which can then exert an influence on HSC activity via cell-cell contact (e.g., cadherin interactions) and secreted biomolecular factors (*22, 23*). Further, the ECM can modulate biotransport of secreted factors via transient binding motifs (e.g. proteoglycans) or steric hindrance to diffusive and advective mass transport (*24-27*). Design of artificial niches offers the opportunity to regulate the transport of biomolecular signals in heterotypic cultures, and engineer HSC-generated autocrine feedback and niche cell paracrine signals. Work by *Pompe et al*. demonstrates this potential with a microcavity system of single vs. doublet HSC cultures, with autocrine feedback dominating quiescent HSC maintenance (*28*).

Expanding upon recent use of exogenous stimuli (e.g. conditioned media) to mimic cell-signaling motifs (*29, 30*), biomaterial design can exploit matrix properties to create regimes of favored cell-signaling mechanism. In a tight matrix, with a small mesh size and low diffusivity, HSCs may experience a small radius of communication, as cell-cell communication is hindered by steric and electrostatic interactions with the matrix, enforcing a mostly autocrine feedback-rich environment. Alternatively, in a loose matrix, with a large mesh size, HSCs have a larger radius of communication, allowing for possible heterotypic paracrine signaling. Culturing hematopoietic lineage positive cells in a three-dimensional collagen system, we recently demonstrated improved maintenance of early progenitor HSC populations in autocrine-dominated environments with low diffusivity (*31*). While promising, this early-generation hydrogel system was not optimal for in-depth studies of the intersection of heterotypic cell interactions and cell mediated remodeling.

Bone marrow mesenchymal stromal cells (MSCs) show significant promise for the development of artificial bone marrow platforms for *ex vivo* HSC culture. MSCs co-localize with HSCs in the perivascular regions and central marrow and influence hematopoietic activity (*32, 33*). Paracrine signaling from proximal MSCs are believed to mediate HSC activation and quiescence via direct cell-cell contact and diffusive signaling (*5, 28, 33-37*) through production of factors such as CXCL12 (*38*), IL-6 (*3, 39*), and TPO (*40*). Biomaterial systems offer the opportunity to identify heterotypic HSC-MSC cultures that support HSC quiescence or self-renewal. However, studies of HSC-MSC interactions are significantly complicated by processes of dynamic reciprocity (*41, 42*). Notably, while the matrix can impart cues to directly influence MSC-HSC interactions, MSCs are also equipped to remodel the matrix via deposition of proteins and enzymatically degradation of the matrix (*41, 43-45*), leading to non-static material properties, and a dynamic feedback loop. MSCs can also create strain gradients and deformation in the local environment, the so-called peri-cellular region (*44*). As a result, it is important to not only consider initial properties of a biomaterial culture system, but to employ methods to interrogate dynamic shifts in these properties, and their effect on resultant domains of signaling. The intricacies of HSC-MSC co-cultures offer unique opportunities to interrogate effects of dynamic reciprocity as a design paradigm in *ex vivo* HSC culture systems.

The objective of this study was to examine the effect of MSC co-culture on HSC quiescence and differentiation patterns within a three-dimensional methacrylamide-functionalized gelatin (GelMA) hydrogel. We employ a library of GelMA hydrogels to examine the integrated influence of hydrogel poroelastic properties, paracrine signals, and MSC-mediated matrix remodeling on HSC lineage specification events. Myeloid lineage specification and progenitor cell quiescent events were examined for murine HSCs cultured in GelMA hydrogels in the presence of increasing concentrations of murine bone marrow MSCs (**Fig. 1**). We also examined HSC differentiation in the context of hydrogel remodeling by MSCs. Together this work represents an integrated approach to consider dynamic biotransport processes as essential design paradigms in biomaterials for the culture and expansion of primary HSC population.

**Figure 1.**
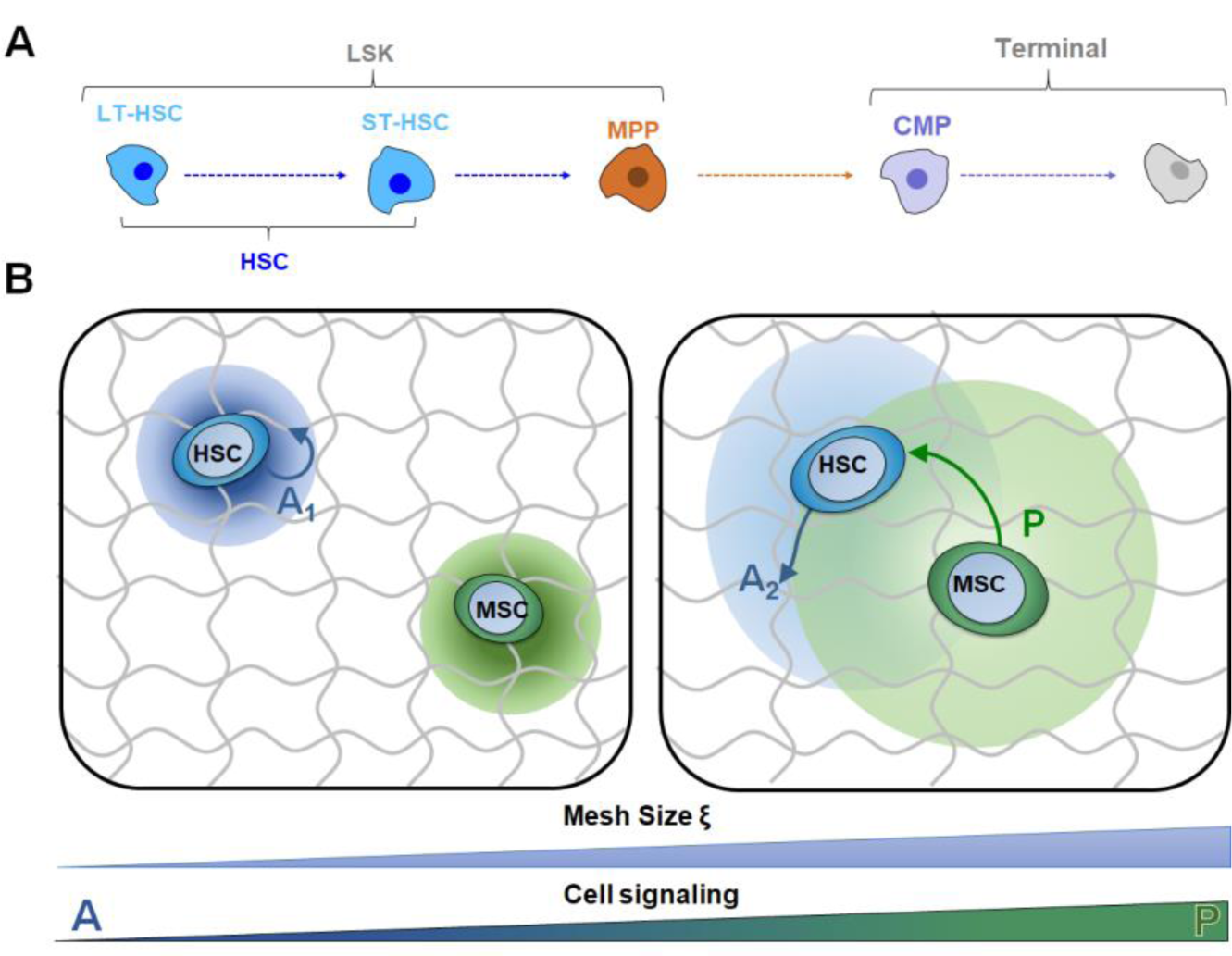
Hematopoietic cell-cell interactions in a dynamic biomaterial landscape. **A)** Hematopoietic lineage hierarchy. Long-term repopulating HSCs and Short-term repopulating HSCs are grouped as a singular HSC population. HSCs and Multipotent Progenitors comprise the HSC progenitor, HSPCs, population. Common Myeloid Progenitors and Lin+ cells make up the Terminal cell population. **B)** Autocrine and paracrine signaling can be tuned by adjusting seeding density and altering biotransport properties. In an HSC-only culture, an increase in mesh size leads to a diffusive regime that increases paracrine signaling. In an HSC-MSC co-culture, small mesh constrains secreted biomolecules and leads to a mostly autocrine signaling. However, as mesh size is increased, autocrine feedback is reduced and paracrine signaling begins to dominate cell-cell interactions.

## Results

### 2.1. Poroelastic characterization of GelMA hydrogels

A library of methacrylamide-functionalized gelatin (GelMA) was assembled and poroelastic parameters were examined in response to changing the degree of methacrylamide functionalization (DOF) of the gelatin macromer, GelMA wt%, and photoinitiator (PI) concentration. The Young’s modulus (E) of the library spanned a range of marrow mimetic values (2.67 ± 1.39 to 32.23 ± 10.7kPa), and showed strong dependence on GelMA wt% and, to a lesser extent, degree of functionalization (DOF) of the GelMA (**Fig. S1**). PI (0.05, 0.1%) was found to only nominally impact the modulus and subsequent analyses here concentrates on the 0.1%PI condition (**Fig. 2A**). The subset of hydrogels fabricated from 4wt% GelMA demonstrated moduli ranging from 4.67 ± 3.20 (35% DOF) to 7.65 ± 3.90kPa (85% DOF). A midrange subset (5wt% GelMA) showed increased elastic moduli range, from 8.90 ± 1.83 (35% DOF) to 13.54 ± 4.81kPa (85% DOF). The highest GelMA content subset (7.5wt%) demonstrated a range of moduli spanning 17.89 ± 10.99 (35% DOF) to 30.06 ± 12.35kPa (35% DOF). Following this analysis, we examined the poroelastic properties of the library of hydrogels in terms of the shear modulus (G) and the diffusion coefficient of water (D_w_). Here, the range of shear modulus spanned 0.36 ± 0.03 to 6.05 ± 0.50kPa (G) while water diffusion ranged from 75.86 ± 11.84 to 211.13 ± 51.84 μm^2^/s (**Fig. S2** and **S3**).

**Figure 2.**
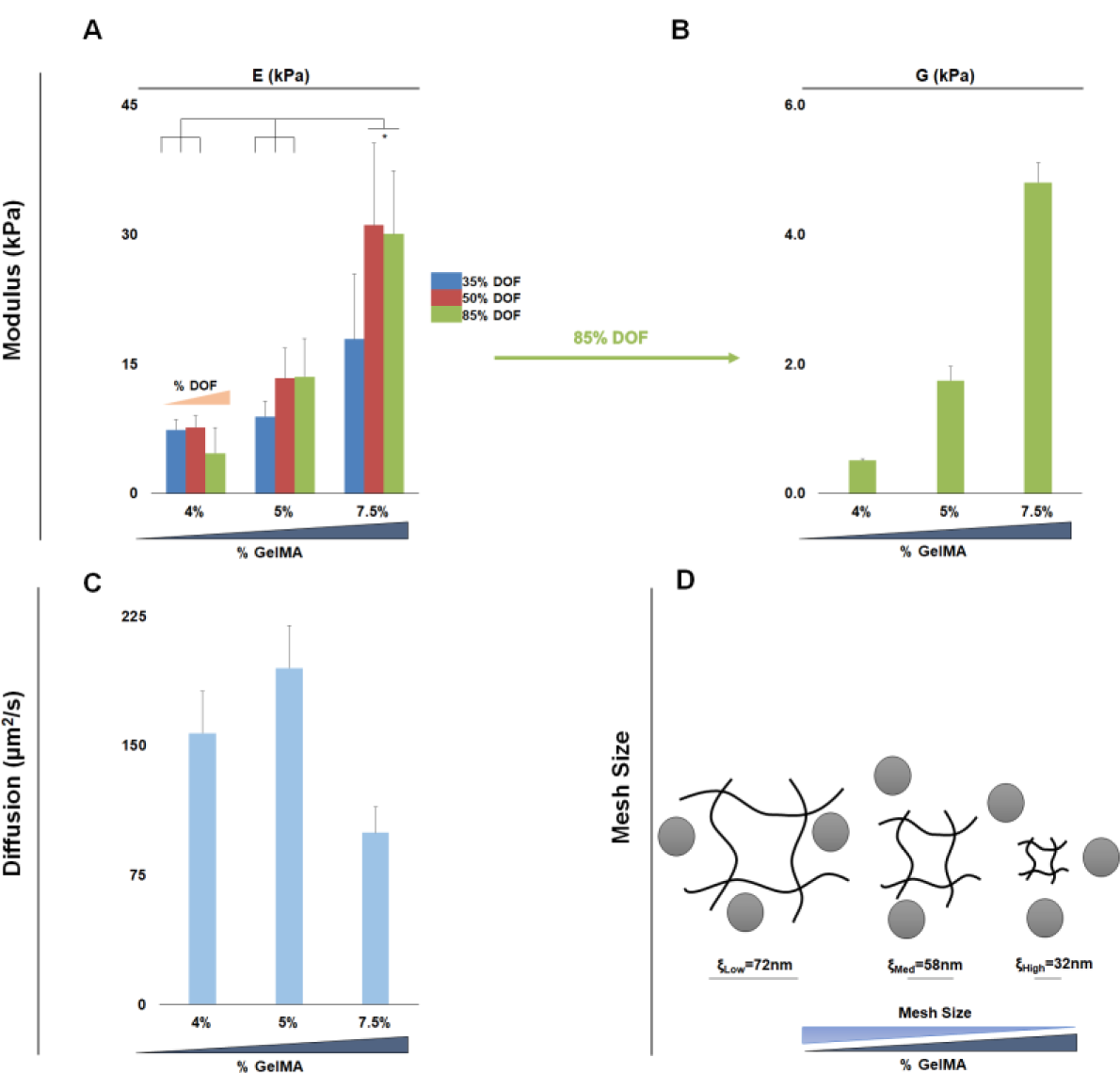
Mechanical characterization of the library of methacrylamide-functionalized hydrogels. (**n = 6**). **A)** Young’s modulus of 9 hydrogel conditions, keeping the PI constant at 0.1% **B)** Shear modulus data from the *Low*, *Med*, and *High* hydrogel variants used for cell culturing, keeping DOF and PI constant at 85% DOF and 0.1%PI. **C)** Diffusion of water in the *Low*, *Med*, and *High* variants exhibits two regimes, *Low* and *Med* (4, 5% GelMA) have a higher diffusion, whereas *High* (7.5% GelMA) has ∼50% reduction in diffusion **D)** Mesh size estimates for the *Low*, *Med*, and *High* variants decreases with increasing GelMA content. There is a trend of *Low* and *Med* exhibiting similar mesh and diffusion, while *High* exhibits a significantly (p < 0.05) lower biotransport regime.

From this complete library of hydrogels, we identified a subset of three hydrogels (85% DOF, 0.1%PI, and 4, 5, 7.5%GelMA) that spanned a range of mechanical properties associated with the native niche: *Low*, E*_Low_* = 4.67 ± 3.20; Med, E*_Med_* = 13.54 ± 4.81; High, E*_High_* = 30.06 ± 12.35kPa (**Fig. 2A**). This subset also displayed a significant increase in the shear modulus (G*_Low_* = 0.51 ± 0.03; G*_Med_* = 1.74 ± 0.22; G*_High_* = 4.80 ± 0.3kPa respectively) (Fig. 2B). Low and Med hydrogels displayed similar water diffusion characteristics (157.19 ± 24.45, 195.04 ± 24.66 μm^2^/s) whereas the High hydrogel variant displayed a significantly reduced D_w_ (99.48 ± 15.11 μm^2^/s) (**Fig. 2C**). Hydrogel mesh size (ξ) calculations showed a mirror inverse of the moduli, with decreased mesh size associated with increased GelMA content: ξ*_Low_*=72, ξ*_Med_*=58, ξ*_High_*=32 nm (**Fig. 2D**).

### 2.2. MSC mediated remodeling of GelMA hydrogels

As MSCs can interact and remodel gelatin-based hydrogels, we quantified MSC remodeling of the *Low*, *Med*, and *High* hydrogels via compressive tests as a function of time in culture (0–7 days) and MSC seeding density (0, 1 × 10^5^, 1 × 10^6^ cells/mL), with results reported as absolute values as well as normalized against the initial properties (day 0) of each variant (**Fig. 3B**). Notably, no observable trends in modulus change were observed over time in either *Low* and *Med* hydrogel variants. However, the *High* hydrogel (7.5%GelMA; 85% DOF; 0.1%PI) displayed significant remodeling over time for the highest MSC seeding density (1 × 10^6^ MSCs/mL). Here, significant biosynthetic remodeling was observed, characterized by significant increases in Young’s modulus (ΔE = 2.61 ± 0.72), almost twice that of other seeding conditions (ΔE = 1.13 ± 0.45, acellular; ΔE = 1.25 ± 0.13, 1 × 10^5^ MSCs/mL) (**Fig. 3A**).

**Figure 3.**
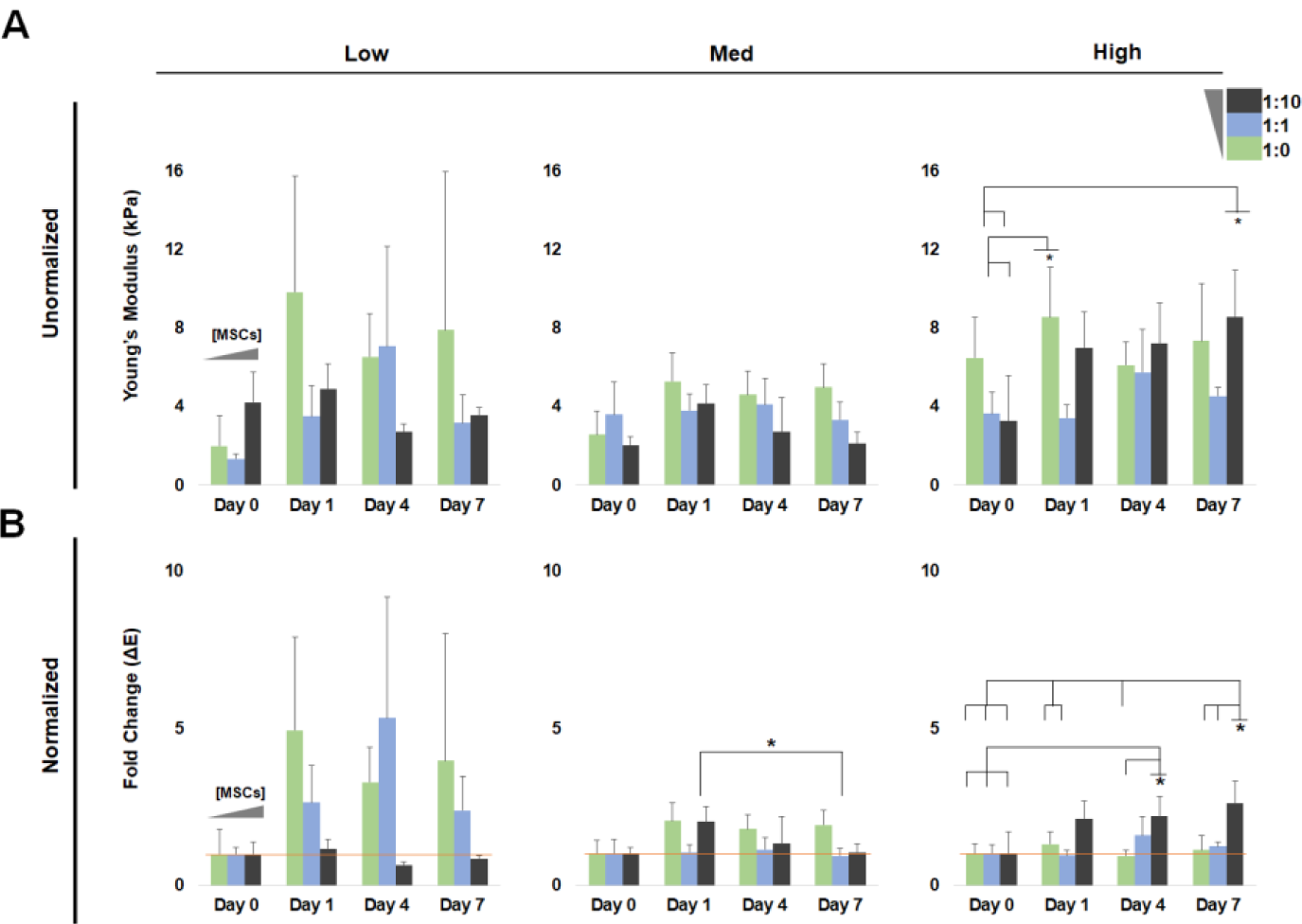
Bulk Young’s modulus of *Low*, *Med*, and *High* hydrogels cultured over 7 days with seeding densities of 0, 1 × 10^5^, and 1 × 10^6^ MSCs/mL (1:0, 1:1, 1:10 HSPCs:MSCs). (**n = 3-6**). **A)** There is minimal change in Young’s modulus over time in any hydrogel and seeding conditions. The high hydrogel shows a significant increase (p < 0.05) from initial seeding to day 7 in the presence of a high concentration of the highest density of MSCs, 1 × 10^6^ MSCs/mL. **B)** The Young’s modulus was normalized to the initial value, day 0, of the specific seeding and hydrogel variant to show fold change. This highlights that only the *High* with the highest concentration of MSCs (1 × 10^6^ MSCs/mL) experiences significant increase (p < 0.05) in bulk Young’s modulus over time.

Expression of remodeling-associated genes overtime was examined to determine local changes in remodeling. PCR shows a downregulation of remodeling associated genes in MSCs as early as day 1, however there were a number of interesting trends across groups. MMP-9 saw the largest upregulation with around 6-fold increase on day 1 across all hydrogel variants in the lowest seeding density (**Fig. 4A**). The high seeding density, 1 × 10^6^ MSCs/mL had the largest MMP-9 upregulation, with 10.8 ± 1.93 fold increase on day 1 in the *Low* hydrogel (**Fig. 4B**). The MMP-activity repressor, tissue inhibitor of metalloprotease, TIMP-1 showed modest upregulation at day 1 (1.3-fold for 1 × 10^5^ MSCs/mL; 1.5-fold for 1 × 10^6^ MSCs/mL) for each hydrogel condition (*Low*, *Med*, *High*), however by intermediate time point, day 4, TIMP-1 had become downregulated. TIMP-2, TIMP-3, and Col1(α1) were sharply downregulated by day 4 hile while MMP-2 showed decreased expression that increased with increasing MSC density 1 × 10^5^ vs. 1 × 10^6^ MSCs/mL) (**Fig. 4A,B**)

**Figure 4.**
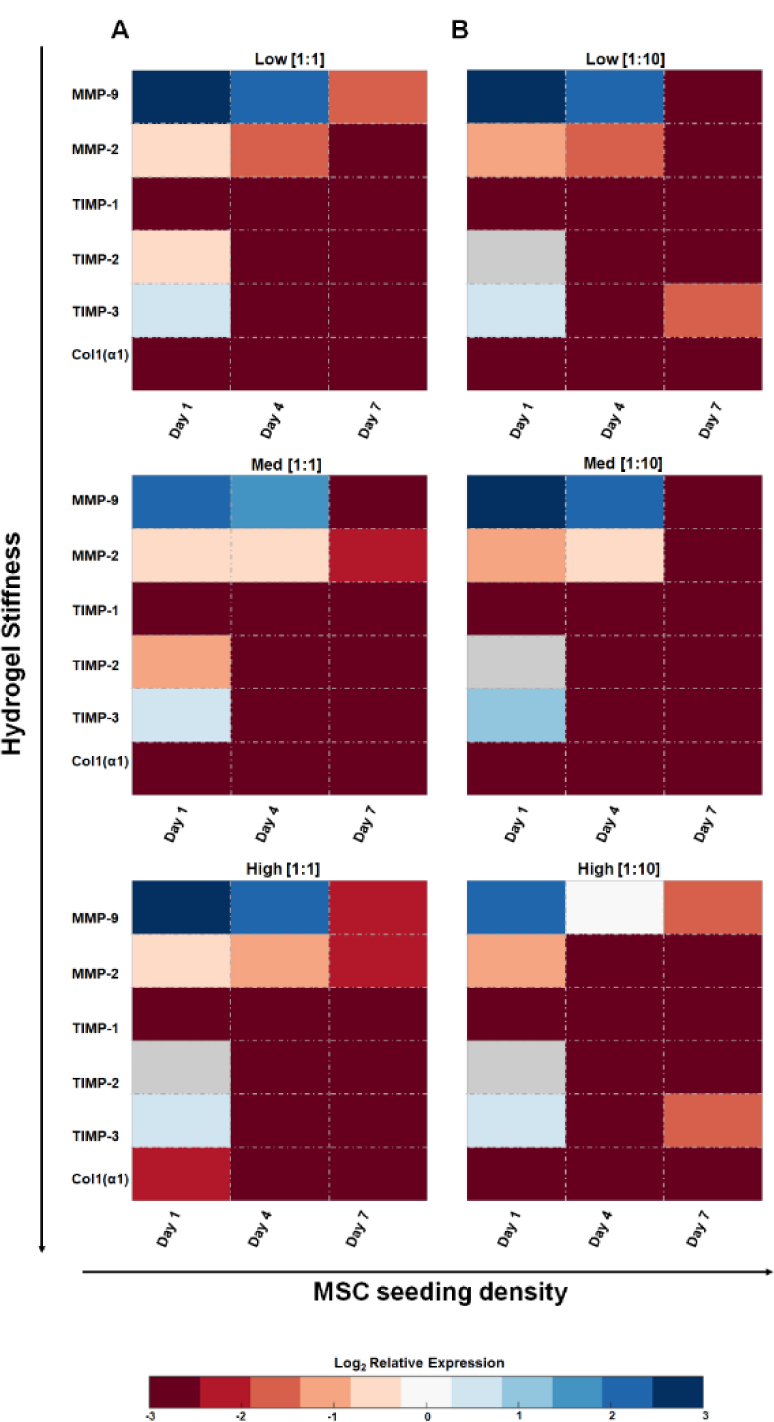
Relative gene expression of matrix-associated genes normalized to day 0 expression, shown on a log2 scale, with downregulation shown in red and upregulation shown in blue. Gray values for TIMP-2 show non-determinant values, indicative of low yields of expression. Matrix degradation genes analyzed were matrix metalloproteases MMP-2 and MMP-9. Matrix additive genes were the tissue inhibitors of metalloproteases TIMP-1, TIMP-2, and TIMP-3, along with the matrix protein collagen, type 1, alpha 1. (**n = 3-6**). **A)** Gene expression of hydrogels seeded with low density of MSCs 1 × 10^5^ MSCs/mL (1:1 HSPCs:MSCs). **B)** Gene expression of hydrogels seeded with high density of MSCs 1 × 10^6^ MSCs/mL (1:10 HSPCs:MSCs).

### 2.3 Culture system dependent hematopoietic differentiation patterns

Murine hematopoietic stem progenitor cells (HSPCs) were cultured in the characterized hydrogel variants at a low seeding density (2000 HSPCs/20 µL solution) to eliminate direct cell-cell contact. Analysis of the homotypic culture displays an HSC population dependence on hydrogel condition: *Low*, *Med*, and *High* (**Fig. 5B,C)**. The percentage of HSCs followed an inverse trend, decreasing with increasing GelMA concentration (**Fig. 5C**). Not unexpectedly, with the decrease in early progenitors, there was an associated increase in the downstream cell population patterns. This was seen in the Common Myeloid Progenitors (CMP) cell population which made up 4.95 ± 1.02% of the hematopoietic cell population in the *Low* hydrogel, but was significantly larger in the *High* hydrogel, 13.72 ± 2.93% (**Fig. 5B**). A similar trend was seen in the terminal cell population, in which increasing GelMA content led to increasing hematopoietic terminal cell population (**Fig. S6**).

**Figure 5.**
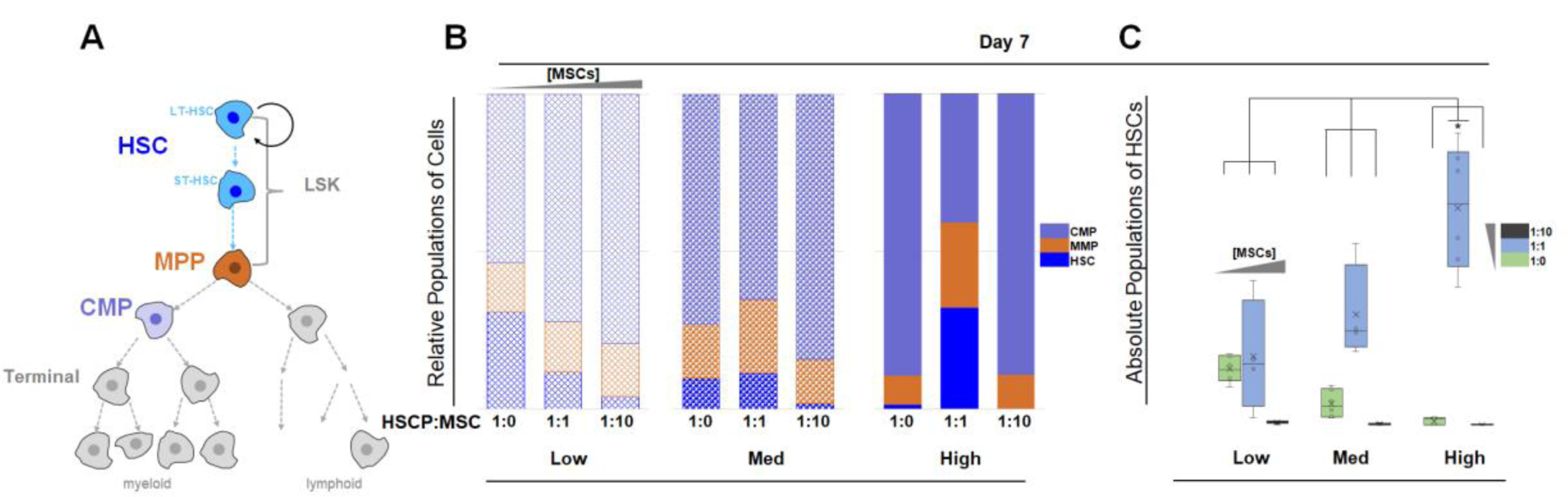
Analysis of cell populations at day 7 (**n = 6**). **A)** The lineage hierarchy of hematopoietic stem cells, with long-term and short-term hematopoietic stem cells grouped as HSCs (blue), Multipotent Progenitors (orange), and Common Myeloid Progenitors (lavender). **B)** The relative population of HSC (blue), MPP (orange), CMP (lavender) are shown for single-culture and co-culture at 1:1 and 1:10 (HSPCs:MSCs), in the *Low, Med*, and *High* variants. The height of each subcolumn reflects how much of the total cell population is made up of the specified cell type. The blue columns (HSC) show that the highest percentage of HSCs are found in the single-culture *Low* variant (1:0), and in the 1:1 co-culture *High* variant. **C)** The percentage of HSCs that make up the total hematopoietic cell population, arranged in order of increasing MSC density (1:0, 1:1, 1:10 HSPCs:MSCs), and in order of increasing hydrogel stiffness (*Low*, *Med*, and *High*). The largest percent of HSC is found in the *High* variant at a 1:1 seeding ratio and is significantly higher (p < 0.05) than all other conditions.

Murine MSCs were also co-cultured with HSPCs in heterotypic cultures with seeding densities kept low to reduce cell-cell contact and ensure only soluble biochemical interactions between cells. These co-culture systems led to important shifts in HSC differentiation patterns (**Fig. 5**). Where in HSC single-cultures the highest final percentage of early progenitor cells was found in the lower GelMA hydrogels, the addition of MSCs in a 1:1 [HSPCs:MSCs] ratio created a reversal in that trend. The highest maintenance was found in the *High* hydrogel variant with HSCs comprising 11.11 ± 3.18% of the hematopoietic population, while the fraction of remaining HSCs was significantly lower in the *Med* and *Low* variants (**Fig. 5B,C)**. From day 1 to day 7 this represents maintenance of 50.2%, 22.08%, and 18.50% of the initial HSPC seeding for *High*, *Med*, and *Low* (**Fig. 6**). Higher HSC maintenance led to a lower terminal cell population of 77.54 ± 5.83% for *High* hydrogel variants (compared to 82.70 ± 6.09% and 91.50 ± 4.67% for *Med* and *Low* hydrogel variants; **Fig. S6**). When HSPCs were co-cultured with a higher density of MSCs (1 × 10^6^ MSCs/mL), at a ratio of 1:10 [HSPCs:MSC], hematopoietic differentiation towards late-stage lineage was enhanced (**Fig. 5B,C)**. Not surprisingly, terminal cells made up the bulk of the hematopoietic population in the 1:10 cultures, representing more than 99% of the hematopoietic cells across all variants (**Fig. S6**).

**Figure 6.**
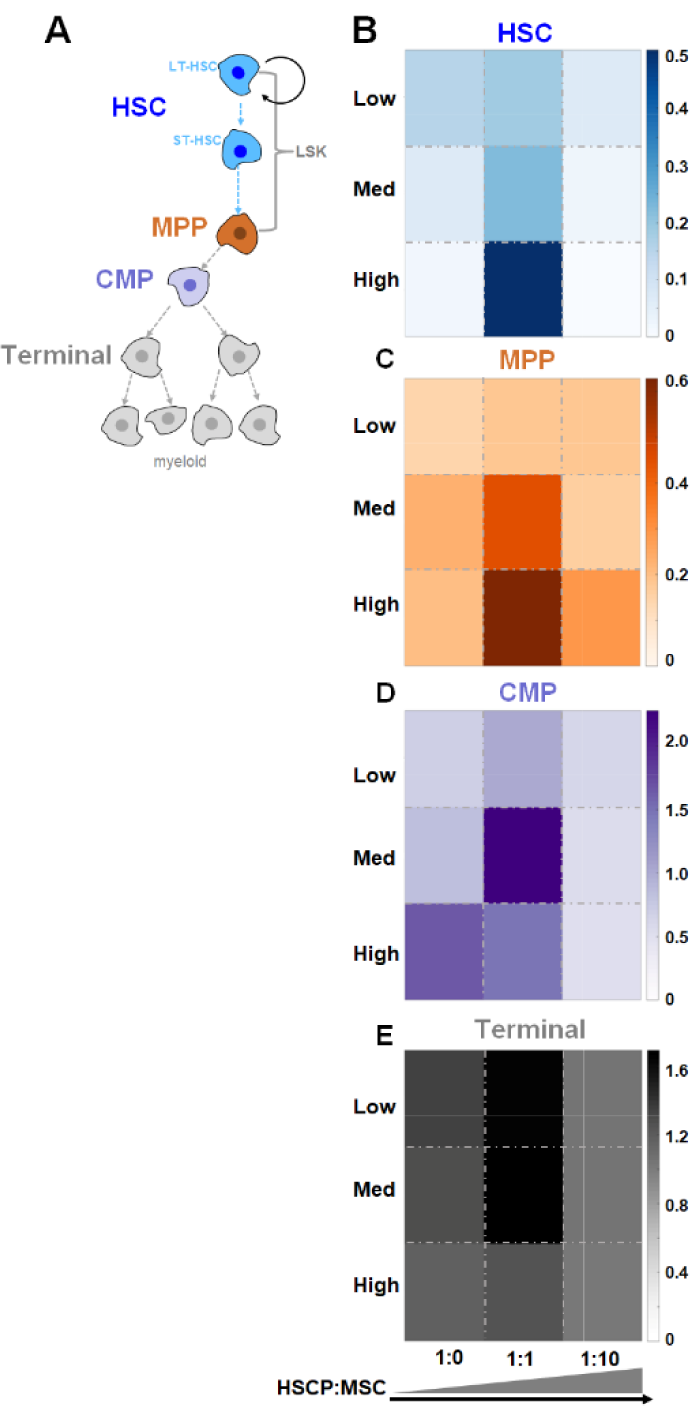
Relative fold change of cell population from day 1 to day 7. **A)** Color-coded hematopoietic lineage hierarchy **B)** Fold change of HSC population. Darker blue indicates higher maintenance of HSC (highest maintenance in *High* variant at 1:1 seeding). **C)** Fold change of MPP population. Darker orange indicates higher maintenance of MPP (highest in *High* variant at 1:1 seeding). **D)** Fold change of CMP population. Darker purple indicates higher maintenance of CMP (highest in *Med* variant at 1:1 seeding). **E)** Fold change of Terminal population. Darker gray indicates higher maintenance of Terminal population (highest in *Med* variant at 1:1 seeding).

### 2.4 Culture dependent shifts in HSC quiescence

We subsequently performed cell cycle analysis for a subset of culture conditions that led to the greatest maintenance of an HSC population when cultured in the absence (*Low* 1:0) vs. presence (*High* 1:1) of MSCs. Here we compare the effect of inclusion of MSCs within the hydrogel environment on maintenance of quiescent HSCs, a key parameter for long-term culture success. Across each cell population and data representation, the inclusion of MSCs in a low diffusive environment (*High* 1:1) led to an increase in the number of quiescent (G0-state) hematopoietic cells. Notably, the inclusion of MSCs lead to a 129-fold (LT-HSC subpopulation) and a 7.7-fold (entire HSC population) increase in the number of quiescent cells versus monocultures of HSCs alone in a highly diffusive environment (*Low* 1:0) (**Fig. 7**). Further, we observed enhanced HSC quiescence as a result of the presence of the hydrogel matrix compared to conventional liquid cultures of HSCs alone or equivalent HSC-MSC mixtures (**Fig. S7**).

**Figure 7.**
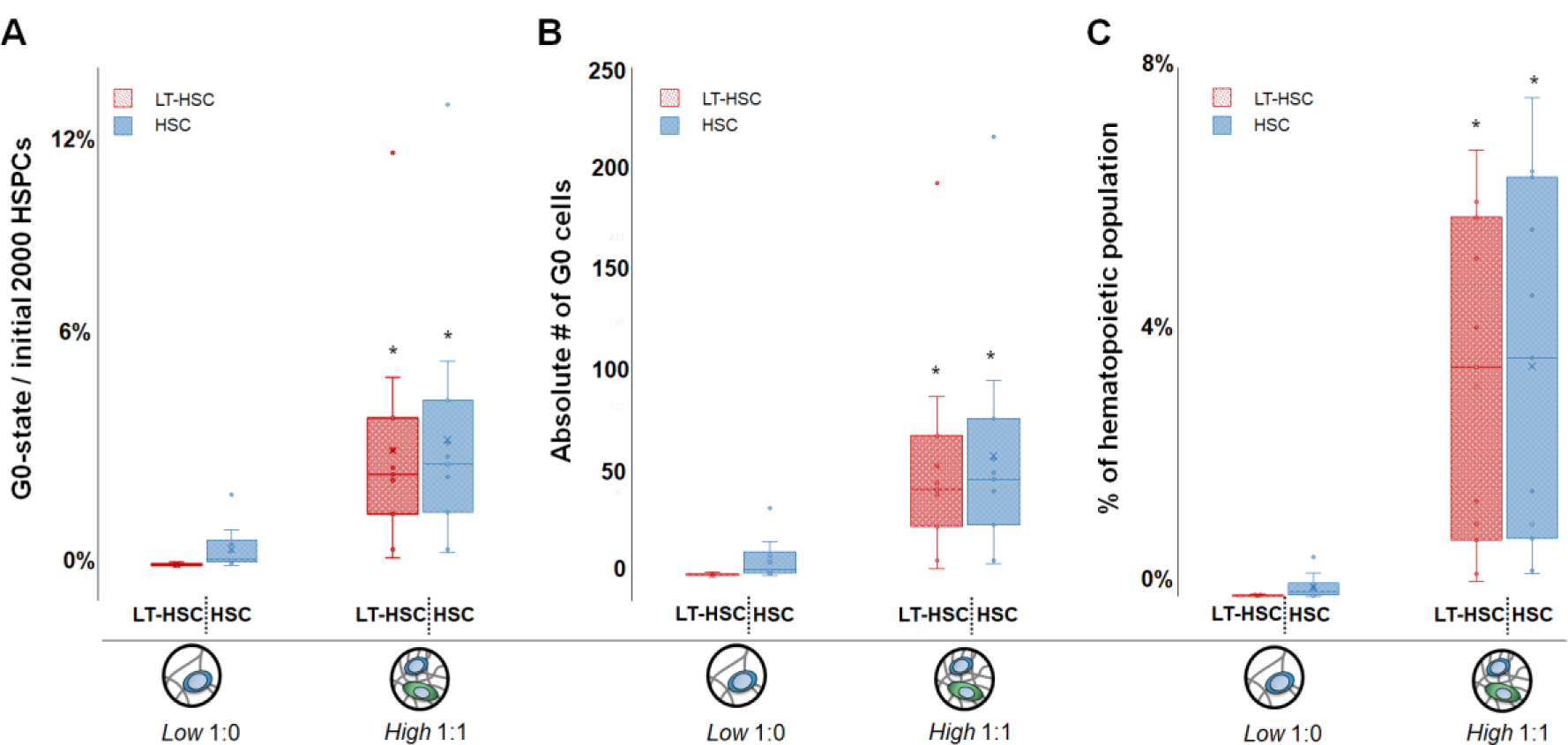
Comparison of LT-HSC and HSC quiescence after 7 days in conditions that led to the greatest maintenance of an HSC population when cultured in the absence (*Low* 1:0 HSPC:MSC) vs. presence (*High* 1:1 HSPC:MSC) of MSCs (**gels = 30, n = 10**). **A)** Fraction of quiescent LT-HSC or HSC, shown as a percentage of the initial 2000 HSPCs seeded per hydrogel condition. **B)** Absolute number of quiescent (G0) LT-HSC and HSCs within each sample. **C)** Percentage of quiescent cells calculated vs. the total number of hematopoietic lineage positive cells after 7 days in culture as a percentage of the overall hematopoietic lineage positive cell population. *: significantly different (p < 0.05) versus the quiescence in high diffusivity hydrogel containing HSCs only (Low 1:0).

## Discussion

Synthetic niche analogs for *ex vivo* culture of HSCs have begun to uncouple the complex signals that are inherent to the native HSC niche. Traditional efforts have demonstrated that changes in matrix composition, mechanics, and selective inclusion of growth factors can substantially alter HSC maintenance and differentiation patterns. Notably, selective interactions with fibronectin (*46-48*), or immobilization of SCF within a hydrogel (*49-51*), can enhance HSC maintenance. More recently, the concept of cell-cell signaling and its role on HSC lineage specification have begun to be examined, primarily in liquid (unhindered-diffusion) experimental systems. *Csaszar et al.* demonstrated that HSC-secreted cytokines can alter *ex vivo* expansion of HSCs within a bioreactor (*52*), while *Müller et al.* demonstrated that individual HSCs cultured in microcavities with autocrine feedback led to increased HSC quiescence (*53*). However, it is also important to consider the influence of heterotypic cell-cell interactions within more complex niche mimics. For this, 3D hydrogel biomaterials are advantageous as heterotypic cultures can be maintained in an environment where the kinetics of HSC-generated autocrine feedback and paracrine signaling from niche-associated cells can be manipulated by the poroelastic properties of the hydrogel itself. Our use of a GelMA hydrogel enables robust control over both mechanical cues and transport of soluble factors to impact HSC lineage and quiescence. Previously, we employed a collagen hydrogel system to examine feedback between HSCs and their progeny of lineage positive cells (*31*). That system demonstrated that signaling modes, i.e. paracrine vs autocrine, in a 3D environment impacted HSC fate decisions, with paracrine signals from Lin^+^ hematopoietic cells capable of enhancing HSC myeloid, while the effects could be tempered in a diffusion-limited hydrogel environment promoted autocrine feedback and HSC maintenance (*31*).

Importantly, while a hydrogel matrix facilitates studies of the influence of mechanical and heterotypic soluble signaling on HSC activity, it may also be sensitive to cell-mediated remodeling. The novelty of this work is the systematic approach to demonstrate the coordinated effects of matrix poroelastic properties, cell mediated remodeling, and heterotypic cell-cell interaction on HSC culture and quiescence. Concepts of dynamic reciprocity, often employed in the context of wound remodeling or the tumor microenvironment (*54, 55*), offer a novel avenue to explore the design of HSC culture platforms. Therefore, we developed a culture system that takes advantage of paracrine signaling and matrix remodeling from niche-associated mesenchymal stromal cells (MSCs). Our application of a library of GelMA hydrogels that span a range of mechanical properties inspired by the native HSC niche (*16*), as well as advanced poroelastic characterization methods (*56, 57*), allows us to examine dynamic relationships between MSCs, the local matrix architecture, and autocrine vs. paracrine domains. HSPCs were co-cultured with MSCs at varying ratios (1:0, 1:1, 1:10 HSPCs:MSCs; 2000 HSPCs) to demonstrate that HSC differentiation patterns can be manipulated by combinations of initial hydrogel properties, MSC mediated remodeling, and the balance of autocrine feedback versus MSC-generated paracrine signals.

Gelatin was chosen as the base material of our system given its inherent ability for cell interaction. It possesses RGD adhesion sites, along with cleavage sites for matrix metalloproteinases produced by MSCs (*58*). Methacrylamide-functionalization and subsequent crosslinking of GelMA reduces the risk of gelatin thermos-instability at elevated temperatures. The mechanical properties of the GelMA hydrogels encompassed the range of moduli observed in the native bone marrow niche (**Fig. 2**). In terms of biotransport, the *Low* and *Med* versus *High* hydrogels displayed two subsets of diffusion and mesh size. Previous results from *Mahadik et al*. (*31*), suggest that the poor diffusivity of the *High* hydrogel is characterized by a small radius of cell-secreted biomolecular diffusion, generating an autocrine-dominated regime for HSC culture (*59*). Similarly, the significantly increased diffusivity of the *Low* and *Med* hydrogels suggests a regime for HSC culture characterized by a large radius of cell-secreted diffusion that reduces the effects of autocrine feedback and enhances MSC-generated paracrine signaling. Dilute cell seeding densities were specifically chosen for this study to promote large cell to cell distances (86-205 µm of spacing between cells, assuming a simple cubic unit). The diffusivity of the gelatin constructs and the dilute seeding conditions (2000 HSPCs/hydrogel) restricted cell-cell interactions to soluble signaling, highlighting the role the hydrogel variants play in determining long-range cell communication.

We observed HSC population maintenance was strongly influenced by initial hydrogel poroelastic properties, the presence of MSCs, and MSC-associated remodeling processes. When cultured in purely homotypic environments, HSC maintenance was greatest in a high diffusion, low stiffness hydrogel (*Low*) (**Fig. 5**). To explore the influence of paracrine signals from niche-associated MSCs, we subsequently examined HSC lineage specification patterns in heterotypic cultures in the same set of *Low*, *Med*, and *High* hydrogels with MSCs at either 1 × 10^5^ or 1 × 10^6^ MSCs/mL (1:1, 1:10 HSPCs:MSCs). The addition of cells did not significantly alter the initial Young’s modulus (**Fig. 3**). The presence of MSCs yielded significant, dynamic matrix remodeling as seen via downregulated matrix deposition related genes (Col1(α1), TIMP-1, TIMP-2, TIMP-3) and upregulated matrix degradation genes (MMP-2, MMP-9) (**Fig. 4**). Together, these results suggest dynamic changes in mesh size and biotransport. While we observed significant matrix remodeling via compressive testing in *High* hydrogels seeded with the highest MSC density, no clear trends were observed in the *Low* or *Med* conditions. Recent results from *Schultz et al.* described non-uniform matrix remodeling (*60*) resulting from differences in diffusive path lengths and reaction times for MMPs vs. TIMPs. This suggests remodeling of our hydrogels was likely non-uniform, making it difficult to resolve local remodeling processes within the heterotypic HSC-MSC cultures with bulk measurements of modulus. However, the significant changes in HSC activity observed as a result of co-culture of MSCs suggest dynamic interaction between HSCs and MSCs is an exciting avenue for further exploration.

The addition of large populations of MSCs (1:10 HSPCs:MSCs) led to a decrease in HSC maintenance for all hydrogel variants. This suggests oversaturation by MSCs likely leads to not only increased soluble signaling but also significantly greater remodeling, both of which reduce maintenance of early hematopoietic progenitors compared to HSC-only cultures (**Fig. 5**). However, addition of an equivalent amount of MSCs (1:1 HSPCs:MSCs) leads to a very different shift in HSC population dynamics, with an overwhelming increase in the ability to maintain an HSC population, most notably an increased fraction of *quiescent* HSCs which is suggested to be protective of long-term hematopoietic homeostasis *in vivo*, and valuable in culture systems for HSC expansion or lineage specification (*61-63*). This increase aligns with literature, confirming the ability of MSCs to influence HSC equilibrium through secreted factors within the *in vivo* niche(*33*). Particularly, the highest maintenance of HSCs in the 1:1 HSPCs:MSCs culture was observed in the *High* hydrogel variant that displayed minimal initial biotransport capabilities but also long-term signatures of MSC-mediated remodeling (MMP-9 expression). This suggests the importance of a balance between initial autocrine-dominated feedback followed by MSC paracrine signals for increasing HSC maintenance. These findings highlight the possibility for selectively employing more complex culture systems to engineer HSC lineage specification patterns.

Bone marrow niches contain a complex milieu of external signals that play a significant role in mediating the activity of HSCs *in vivo*. The development of a synthetic niche analog for HSC culture can take advantage of poroelastic properties to mediate the balance of heterotypic cell signaling via soluble factors. Here, we explored the use of MSC-secreted paracrine signaling as well as MSC-mediated matrix remodeling as a means to increase HSC maintenance during *in vitro* culture using a library of GelMA hydrogels. Examining HSC differentiation and cell-cycle patterns, as well as dynamic remodeling of a model GelMA hydrogel in the presence of MSCs suggests a path towards *ex vivo* expansion of HSCs. We show that combinations of initial hydrogel properties and a moderate amount of MSCs increases the maintenance of HSCs in a culture system with initially hindered diffusion. These findings suggest that MSC-secreted paracrine signaling and initial autocrine feedback are essential *in vitro* tools seeking to expand HSC populations. Such findings offer exciting opportunities to explore concepts of dynamic reciprocity in the context of heterotypic cell cultures. Matrix remodeling as well as selective domains of autocrine feedback versus paracrine signaling can be leveraged for the intelligent design of an *in vitro* HSC culture systems.

## Materials and Methods

### 4.1 Material characterization

#### 4.1.1 GelMA synthesis and hydrogel fabrication

Porcine Type A gelatin (Sigma, St. Louis, MO) was functionalized along its backbone with pendulant methacrylamide groups following a previously described protocol (*58, 64*). In brief, methacrylate anhydride (MA) was added dropwise to a solution of gelatin in PBS before quenching in excess PBS, purification via dialysis, and lyophilization. The ratio of gelatin to MA was tuned to control the degree of functionalization (35%, 50%, 85% DOF; quantified by H^1^ NMR) (*58*). Methacrylamide-functionalized gelatin (GelMA) hydrogel variants were subsequently formed from 20 µL of GelMA precursor suspension mixed with a lithium acylphosphinate (LAP) photoinitiator (PI) in circular Teflon molds (5 mm dia.) exposed to 7.14 mW/cm^2^ UV light for 30 seconds. After crosslinking, hydrogels were immediately hydrated in PBS (*49, 65*).

#### 4.1.2 Determination of poroelastic properties

A library of 18 acellular hydrogels were fabricated from 35, 50, and 85% DOF GelMA at 4, 5, and 7.5% (w/v) GelMA, with 0.05 or 0.1% (w/v) PI. The Young’s modulus was determined using an Instron 5943 mechanical tester in unconfined compression (Instron, Norwood, MA). Briefly, a 0.005N preload was applied followed by compression at a rate of 0.1 mm/min until 20% compression was reached. The Young’s modulus was taken as the slope of the linear fit applied to 10% of the stress vs. strain data using Origin Statistical Software (Northampton, MA) (*66, 67*). An offset of 0 or 5% strain was applied prior to linear fit analysis to confirm that the compression platen was completely in contact with the hydrogel surface. The elastic moduli of cellular hydrogels containing 1 × 10^5^ or 1 × 10^6^ MSCs/mL (experimental description below) were traced over seven days in culture, using the same compression protocol. After mechanical analysis, MSC-hydrogels were immediately put on dry ice and stored at −80°C for subsequent gene expression analyses.

Stress-relaxation indentation tests were performed on acellular hydrogels using MFP-3D AFM (Asylum Research, Goleta, CA), with a 4.5 µm spherical tip (Novascan, Ames, IA). The generated force curve was used to extract out the shear modulus and diffusion coefficient of water. Specimens were rapidly indented to 0.5, 1, or 2 µm, and held for a test time of 5 seconds. This process has been described elsewhere, but in brief, the instantaneous force was normalized to the far-field force and fitted to a finite element analysis formula by modulating the characteristic time and material properties (shear modulus, diffusion) (*56, 57, 68, 69*).

#### 4.1.3 Mesh size estimate

The mesh size of the hydrogel library was determined using hydrogel swelling ratios and is a function of the volume fraction and the mean-squared end-to-end distance of the gelatin chain (*70-72*).

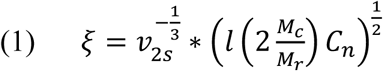

In determining material parameters, it was assumed that the gelatin chains were long enough to neglect chain-end effects. It was also assumed that the molecular weight of the repeat unit, 𝑀_𝑟_, was the averaged molecular weight of the amino acid composition. Amino Acid Analysis was performed at the UC Davis Genome Center with a Hitachi L-8900 (Li-based) analyzer. The mass swelling ratio was determined by the ratio of the dry to wet weight. The swelling ratio was determined immediately following crosslinking or after 24hrs in PBS, respectively known as relaxed and equilibrium states. Hydrogels were weighed before and after lyophilization to obtain the mass swelling ratio, and then transformed to volumetric swelling ratio following the equation below, using the solvent density (𝜌𝑃𝐵𝑆 = 1.01𝑔/𝑐𝑚^3^) (*73*) and gelatin density (𝜌𝑔𝑒𝑙𝑎𝑡𝑖𝑛 = 1.345𝑔/𝑐𝑚^3^) (*74-76*).

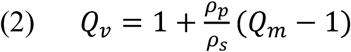

From this, the volume fraction of gelatin is calculated as:

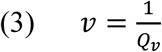

The molecular weight between crosslinks, 𝑀_𝑐_, was estimated using the well-known Flory-Rehner equation adapted by Bray and Merrill for use in polymers crosslinked in solvent (*71, 77*). The applicability of the modified equation for specific concentration regimes was confirmed by Peppas and Merrill (*70*). The 𝑀_𝑐_ is a function of material properties such as the molar weight of the solvent, 𝑉_1_, the specific volume of the polymer, 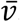, and the solvent-polymer interaction, 𝑋_1_, also known as Flory’s Chi parameter. It is also a function of the system parameters such as the volume fraction of the polymer in its relaxed state and in the equilibrium state, 𝑣_2𝑟_ and 𝑣_2𝑠_, and the number average molecular weight before crosslinking, 𝑀_𝑛_.

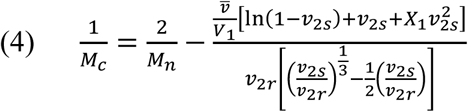

Flory’s characteristic ratio,𝐶_𝑛_, can be approximated by a constant number as the number of monomers goes to infinity, 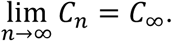 For a stiff chain, this can be approximated using the worm-like or Kratky-Porod persistence model by using the persistence length, 𝑙_𝑝_, and the monomer unit length, and modified for proteins (*78, 79*):

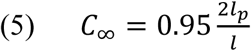

### 4.2 Hematopoietic stem cell isolation

All work involving primary cell extraction was conducted under approved animal welfare protocols (Institutional Animal Care and Use Committee, University of Illinois at Urbana-Champaign). Primary murine hematopoietic stem progenitor cells (HSPC) were extracted from the femur and tibia bone marrow of C57BL/6 female mice, age 4 – 8 weeks (Jackson Labs). The bones were gently crushed with pestle and mortar and washed with PBS + 5%FBS prior to filtration with a 40 µm sterile filter (*31, 49*). The collected suspension was lysed with ACK lysis buffer (Invitrogen, Carlsbad, CA), and then rinsed with PBS/FBS. Initial HSPC isolation was performed using defined protocols of EasySep™ Mouse Hematopoietic Progenitor Cell Enrichment Kit and Magnet (#19756, #18000, Stemcell Technologies, CA). A secondary isolation further enriched the HSPC population by collecting the Lin^−^ Sca-1^+^ c-kit^+^ (LSK) fraction using a BD FACS Aria II flow cytometer. All antibodies were supplied by eBioscience (San Diego, CA), and are as follows: PE-conjugated Sca-1 (1:100 dilution), APC-efluor780-conjugated c-kit (1:100 dilution), and a cocktail of FITC-conjugated lineage specific surface markers (CD5, B220, Mac-1, CD8a, Gr-1, and Ter-119) (*80-82*). The LSK faction was collect in PBS + 25% FBS on ice and immediately used (*13, 21, 31, 49, 83-85*).

### 43 Mesenchymal stromal cell isolation

Murine MSCs were isolated from the femur and tibia bones used for HSPC extraction. Following gentle crushing with a pestle and mortar, MSCs were isolated from bone following commercially available defined protocols of MesenCult™ Proliferation Kit with MesenPure™ (#05512 Stemcell Technologies, CA). The MSCs were cultured in one T25 flask per mouse and allowed to expand for two weeks prior to collection (zero passage), with a half-media change after one week. Non-adherent cells were washed away, and the adherent cells were collected after trypsinization with Gibco TrypLE (ThermoFisher Scientific, Waltham, MA). Cells were collected and immediately used.

### 4.4 Hematopoietic stem cell culture in GelMA hydrogel in the presence of MSCs

Single culture HSPCs were encapsulated in GelMA hydrogels at a density of 1 × 10^5^ HSPCs/mL. Similarly, co-cultures of HSPCs and MSCs were encapsulated at a ratio of 1:1 or 1:10, HSPCs to MSC, keeping a constant density of 1 × 10^5^ HSPCs/mL. HSPCs or MSCs in a suspension were added to the hydrogel precursor solution at the desired seeding density of 1 × 10^5^ HSPCs/mL and 1 × 10^5^ or 1 × 10^6^ MSCs/mL, while maintaining the concentration of GelMA. Upon crosslinking, each hydrogel-cell construct was placed into an individual well of a 48-well plate, and cultured at 37°C (5% CO_2_) for 7 days in 300 µL StemSpan™ SFEM (#09650 Stemcell Technologies) supplemented with 100 ng/mL SCF (Peprotech) and 0.1% Pen Strep (Gibco), with media changes every 2 days (*21*).

### 4.5. Analysis of HSC differentiation patterns

Differentiation patterns of HSPCs were analyzed via Fluorescence-Assisted Cytometry (FACs), using a BD LSR Fortessa (BD Biosciences, San Jose, CA). Cell seeded hydrogels were degraded with 100 Units of Collagenase Type IV (Worthington Biochemical, Lakewood, NJ) for 20 minutes at 37°C, and gently mixed via pipetting every 10 minutes. The reaction was quenched, and the cells were re-suspended in 100 µL PBS and 5% FBS and stained with a cocktail of antibodies (eBioscience, San Diego, CA) to profile HSC lineage specification: eFluor 660 CD34 (1:100 dilution, 90 mins), PerCP-eFluor 710 CD135 (1:100 dilution, 30 mins), eFluor 450 CD16/CD32 (1:200 dilution, 30 mins), APC-eFluor 780 c-Kit (1:800 dilution, 30 mins), PE Sca-1 (1:400 dilution, 30 mins), FITC lineage cocktail (CD5, B220, Mac-1, CD8z, Gr-1, Ter-119, 1:100 dilution, 30 min). Hematopoietic cells were subsequently classified as more primitive Long-Term repopulating HSCs (LT-HSCs: CD34^lo^ CD135^lo^ Lin^−^ Sca1^+^ c-kit^+^) (*84, 86, 87*); differentiated Short-Term repopulating HSCs (ST-HSCs: CD34^hi^ CD135^lo^ LSK) (*84, 86, 87*); more differentiated Multipotent progenitors (MPPs: CD34^+^ CD135^+^ LSK) (*87, 88*), and lineage-specified Common Myeloid Progenitors (CMPs: Lin^−^ c-kit^+^ Sca1^−^ CD34^+^ CD16/32^−^) (*88*), (*81, 89-91*). Individual cell populations are reported as percentages of the total hematopoietic cell population in each culture, with terminal cells classified as hematopoietic cells that were not part of the LSK fraction. For some cell population analyses, LT-HSC and ST-HSC were grouped together as a common HSC population.

### 4.6. Analysis of cell cycle state

Conditions identified as optimal in maintaining HSC numbers (HSC only in *Low* hydrogel variant; 1:1 HSC:MSC co-culture in *High* hydrogel variant) were chosen for subsequent cell cycle analysis. The 1:0 *Low* and 1:1 *High* (HSPC:MSC) conditions were cultured and cells were isolated at day 7 as described above. Experimental replicates (n = 10/condition) were performed; each sample contained three separately cultured hydrogels (n_gels_=30). Isolated cells were then live/dead stained with LIVE/DEAD™ Fixable Red Dead Cell Stain Kit (#L34971, ThermoFisher) following the standard protocol, and LT-HSC, ST-HSC, and MPP populations labeled via antibody cocktail (defined above). Stained cells were fixed and permeabilized with Foxp3 / Transcription Factor Staining Buffer Set (#00-5523-00, eBioscience), and stained with BV605-conjugated Ki-67 (1:100 dilution) and DAPI (#D21490, ThermoFisher). Quiescent cells (DAPI^2n^ Ki-67^−^) from each hematopoietic population were subsequently analyzed via Flow Cytometry.

### 4.7. Analysis of MSC mediated-remodeling profiles

Matrix remodeling associated gene expression profiles were examined for MSC-seeded hydrogel specimens via QuantiStudio 7 Flex real-time PCR (Applied Biosystems). Frozen hydrogels isolated at days 0, 1, 4, and 7 during culture were crushed using a pestle, with mRNA extracted using RNeasy Plant Mini Kit (Qiagen, Germany). mRNA to cDNA was accomplished using the QuantiTect Rev. Transcription kit (Qiagen) and prepped for RT-PCR using PIPETMAX (Gilson, Middleton, WI). The CT of each gene/sample was performed in duplicate. Results were analyzed using the comparative CT method (2^−ΔΔCT^). Primer specificity and efficiency were validated prior to comparative CT analysis (92). Matrix deposition genes examined were collagen, type I, alpha 1 (Col1a1), and tissue inhibitors of metalloprotease TIMP-1, TIMP-2, and TIMP-3. Matrix degradation genes examined were matrix metalloproteases MMP-2 and MMP-9, also known as Gelatinase-A and Gelatinase-B (*93*). Expression of glyceraldehyde-3-phosphate dehydrogenase (GAPDH) was used for housekeeping. Gene expression was first normalized to the housekeeping gene (GAPDH), and then compared against the gene expression at day 0 of each specific seeding and hydrogel condition (**Fig. S4**). Primer information can be found in **Table S1.**

### 4.8. Statistics

All statistics were performed on a minimum of 3 replicates. Significance was tested with Origin Statistical Software using 2-way ANOVA, at a significance level of 0.05. Normality of the data was determined using the Shapiro-Wilkes test at significance 0.05 (*94*), and homoscedasticity (equality of variance) determined using Levene’s Test (*95*). Significance of quiescence was determined with the nonparametric Mann-Whitney test (2-sample) at the 0.05 significance level. Post-hoc power analysis was used to confirm the sample size was large enough to maintain a maximum Type I error of *α*=0.05, and Type II error of *β*=0.20 (*96, 97*). Young’s modulus outliers were removed using Dixon’s Q-test at significance level 0.05 (*98*). Means are reported with their associated standard deviations. In the comparative C_T_ method of gene expression, the error was propagated throughout the calculations.

## Supporting information

supplemental only

## Acknowledgments

The authors would like to acknowledge Dr. Barbara Pilas of the Roy J. Carver Biotechnology Center (Flow Cytometry Facility, UIUC) as well as Dr. Bhushan Mahadik and Dr. Ji-Sun Choi (ChBE, UIUC) for assistance with bone marrow cell isolation and flow cytometry. Preliminary Young’s modulus testing and mRNA extraction was aided by undergraduate researchers Kirsten Schroeder and Gabrielle Wolter respectively. Research reported in this publication was supported by the National Institute of Diabetes and Digestive and Kidney Diseases of the National Institutes of Health under Award Numbers R01 DK099528 (B.A.C.H) and F31 DK117514 (A.E.G.), as well as by the National Institute of Biomedical Imaging and Bioengineering of the National Institutes of Health under Award Numbers R21 EB018481 (B.A.C.H.) and T32 EB019944 (A.E.G.). The content is solely the responsibility of the authors and does not necessarily represent the official views of the NIH. Amino Acid Analysis was performed at the Molecular Structure Facility in the UC Davis Genome Center at the University of California, Davis. Material characterization was carried out in part in the Frederick Seitz Materials Research Laboratory Central Research Facilities, University of Illinois. The authors are also grateful for additional funding provided by the Department of Chemical & Biomolecular Engineering and the Institute for Genomic Biology at the University of Illinois at Urbana-Champaign.

## Supplementary Materials

**Supplemental Figure 1.**
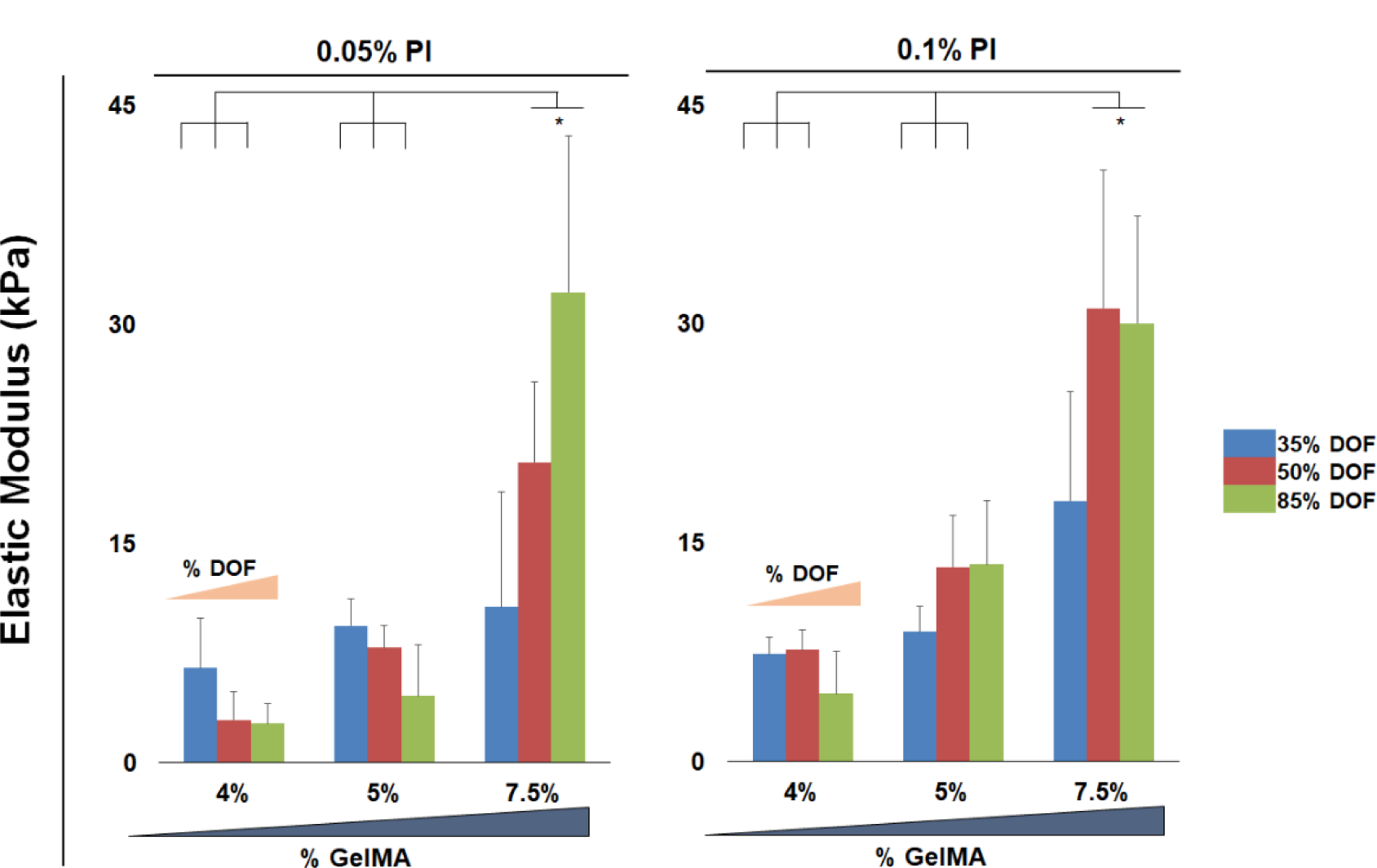
Compressive mechanical analysis of the complete library of hydrogels. The degree of methacrylamide-functionalization was altered from 35, 50, 85%, and the content of GelMA was increased from 4, 5, 7.5%. * marks significantly different at the p < 0.05 level. (**n = 6**). **A**) Elastic modulus of hydrogels prepared with 0.05% photoinitiator. **B**) Elastic modulus of hydrogels prepared with 0.1% photoinitiator.

**Supplemental Figure 2.**
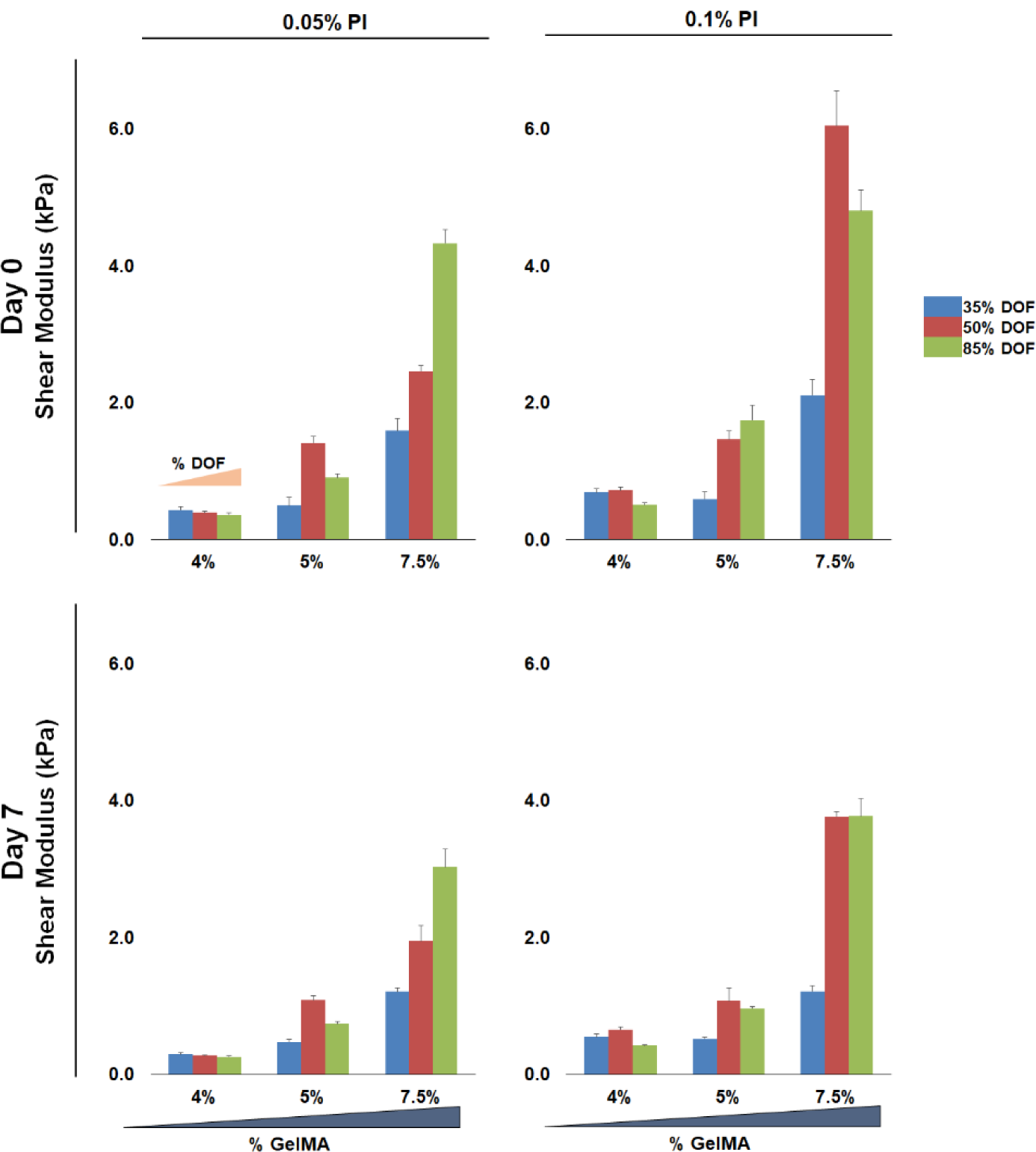
Shear modulus characterization of the complete library of hydrogels. The degree of methacrylamide-functionalization was altered from 35, 50, 85%, and the content of GelMA was increased from 4, 5, 7.5%. (**n = 6**). **A**) Shear modulus of hydrogels prepared with 0.05% photoinitiator, measured at Day 0 after hydration in PBS. **B**) Shear modulus of hydrogels prepared with 0.1% photoinitiator, measured at Day 0 after hydration in PBS. **C**) Shear modulus of hydrogels prepared with 0.05% photoinitiator, measured at Day 7. **D**) Shear modulus of hydrogels prepared with 0.1% photoinitiator, measured at Day 7.

**Supplemental Figure 3.**
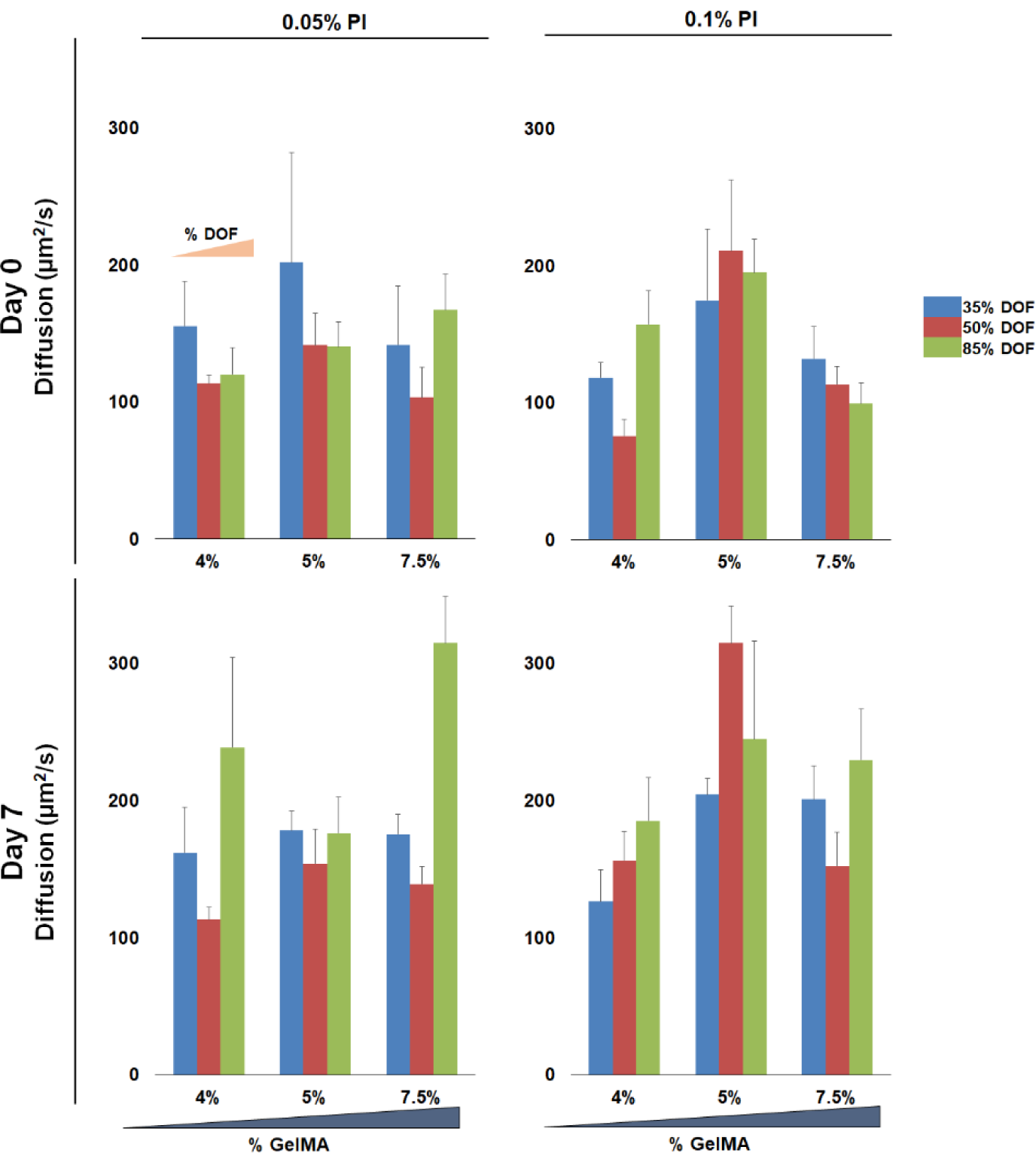
The diffusion coefficient of water in each hydrogel was measured for the complete library of hydrogels. The degree of methacrylamide-functionalization was altered from 35, 50, 85%, and the content of GelMA was increased from 4, 5, 7.5%. (**n = 6**). **A**) Diffusion coefficient of water in hydrogels prepared with 0.05% photoinitiator, measured at Day 0 after hydration in PBS. **B**) Diffusion coefficient of water in hydrogels prepared with 0.1% photoinitiator, measured at Day 0 after hydration in PBS. **C**) Diffusion coefficient of water in hydrogels prepared with 0.05% photoinitiator, measured at Day 7. **D**) Diffusion coefficient of water in hydrogels prepared with 0.1% photoinitiator, measured at Day 7.

**Supplemental Figure 4.**
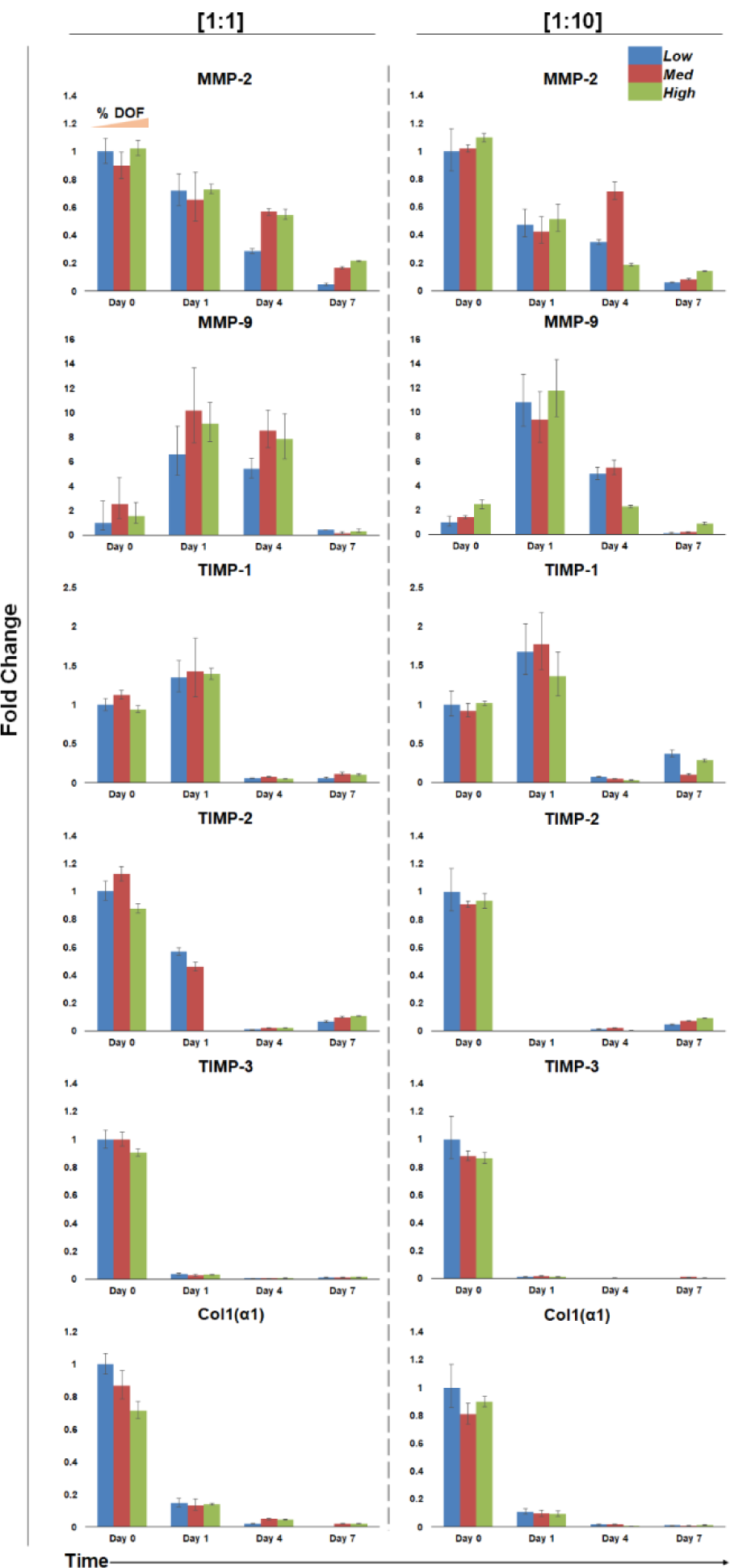
Relative expression of matrix-associated genes in *Low*, *Med*, and *High* hydrogels seeded with 1 × 10^5^ (1:1) and 1 × 10^6^ MSCs/mL (1:10). GAPDH was the housekeeping gene, and expression was normalized to Day 0 of the low hydrogel condition. (**n = 3-6**). **A**) Relative gene expression of hydrogels seeded with 1 × 10^5^ MSCs/mL (1:1) **B**) Relative gene expression of hydrogels seeded with 1 × 10^6^ MSCs/mL (1:10).

**Supplemental Figure 5.**
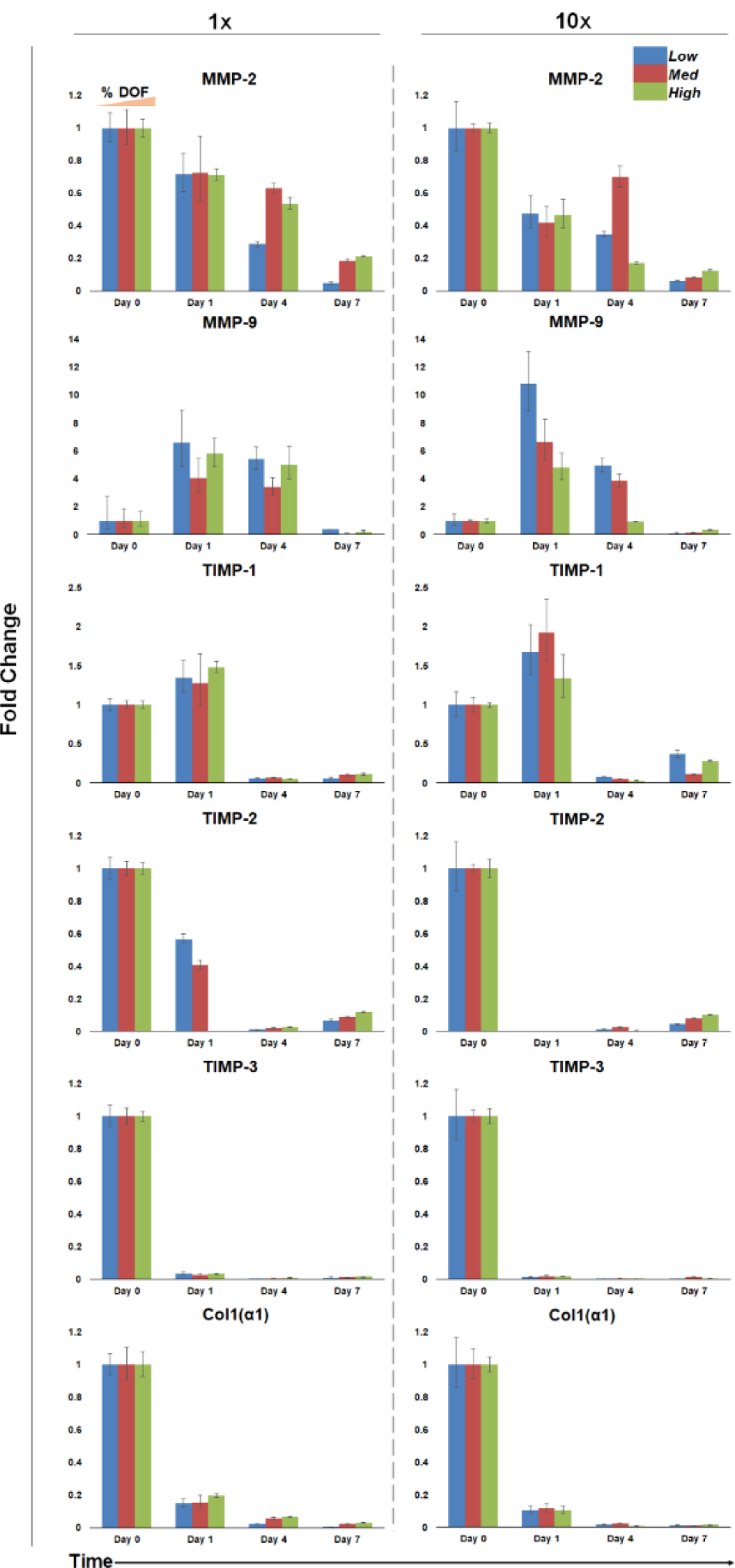
Relative expression of matrix-associated genes in *Low*, *Med*, and *High* hydrogels seeded with 1 × 10^5^ (1:1) and 1 × 10^6^ MSCs/mL (1:10). GAPDH was the housekeeping gene, and expression was normalized to Day 0 of the specific hydrogel and seeding condition. (**n = 3-6**). **A**) Relative gene expression of hydrogels seeded with 1 × 10^5^ MSCs/mL (1:1). **B**) Relative gene expression of hydrogels seeded with 1 × 10^6^ MSCs/mL (1:10).

**Supplemental Figure 6.**
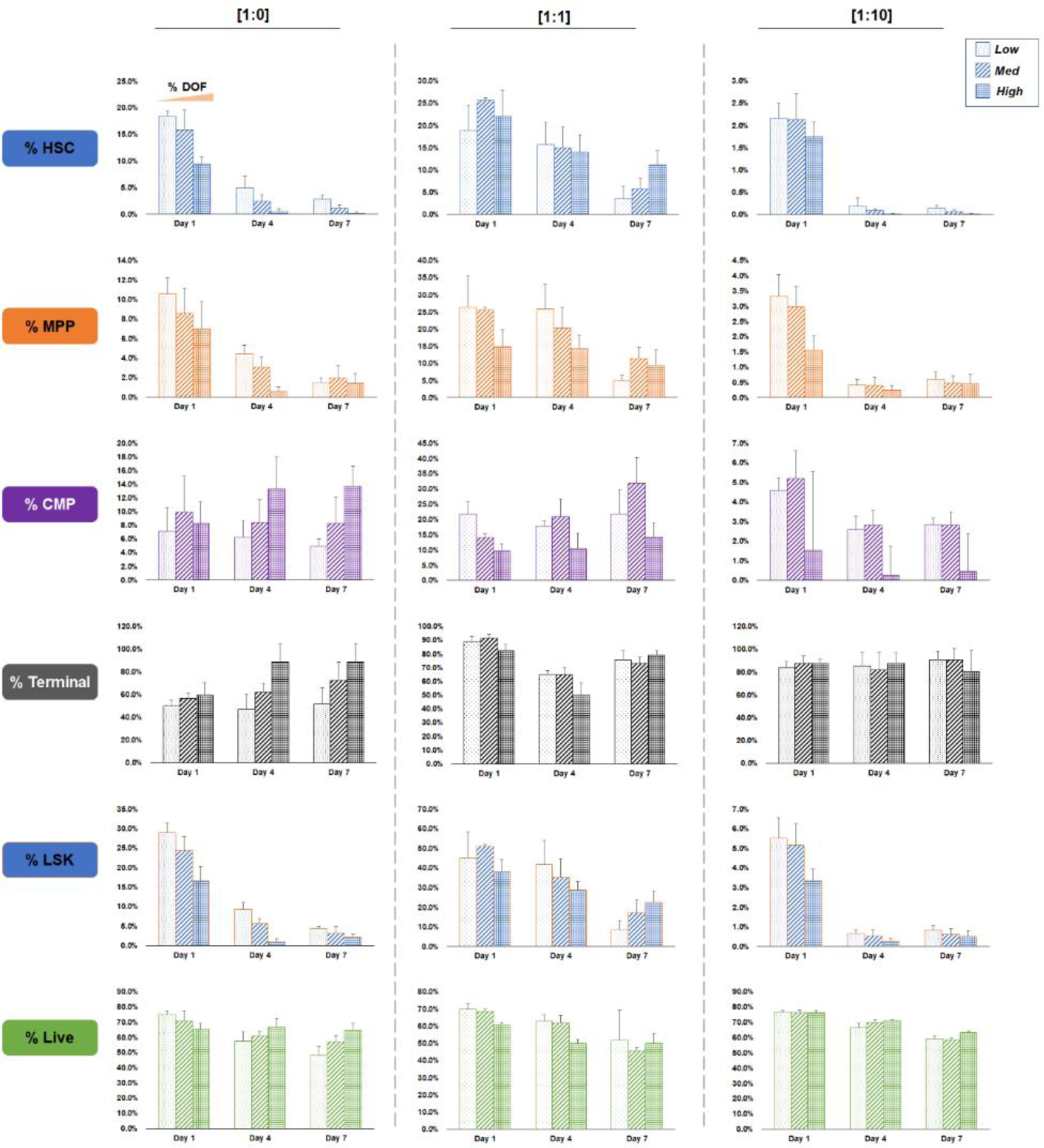
Hematopoietic cell differentiation patterns within *Low*, *Med*, and *High* hydrogels with 0, 1 × 10^5^, and 1 × 10^6^ MSCs/mL (1:0, 1:1, and 1:10 respectively). Each cell type is expressed as a percentage of the overall hematopoietic cell population. (**n = 6**). **A**) Hematopoietic stem cell population, comprised of long-term and short-term HSCs. **B**) Multipotent progenitor cells. **C**) Common myeloid progenitor cells. **D**) Terminal cell population, comprised of all non-LSK hematopoietic cells. **E**) LSK cells comprised of LT-HSC, ST-HSCs, and MPPs. **F**) The live population of cells expressed as a percentage of live cells over the total population of all cells.

**Supplemental Figure 7.**
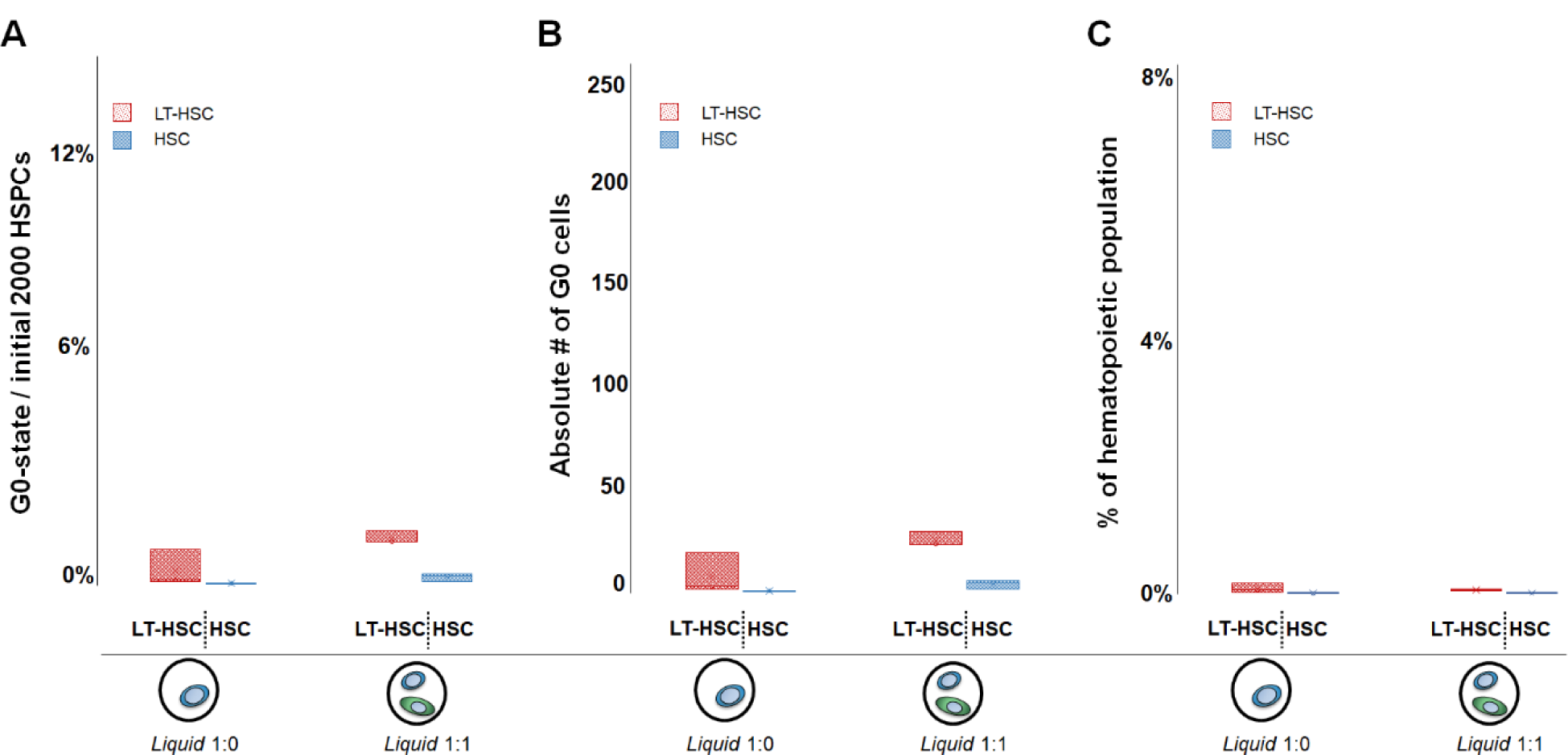
Number and fraction of quiescent LT-HSCs and HSCs in liquid culture containing either 1 × 10^5^ HSPCs/mL alone (1:0), or a mixture of 1 × 10^5^ HSPCs/mL and 1 × 10^5^ MSCs/mL (1:1). (n = 3). **A)** Number of quiescent LT-HSC or HSC, shown as a percentage of the initial 2000 HSPCs seeded. **B)** The absolute number of quiescent (G0) LT-HSC and HSCs within each sample (both populations are highest in the *High* variant at 1:1 seeding). **C)** Percentage of quiescent cells calculated vs. the total number of hematopoietic lineage positive cells after 7 days in culture (including hematopoietic progeny that arise during the culture period).

**Supplemental Table 1.**
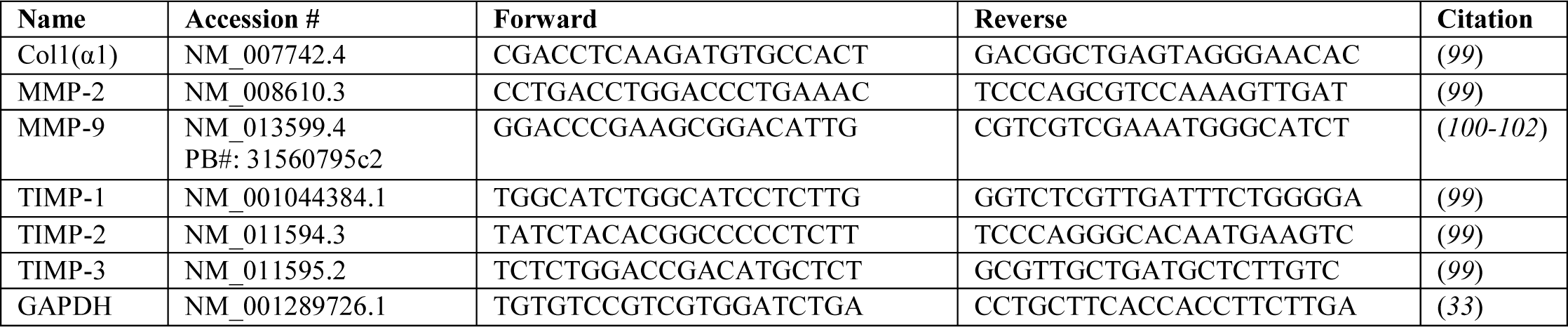
Forward and reverse primer sequences for matrix associated genes. Sequences were made using NCBI Primer-blast or taken from literature.

**Supplemental Table 2.**
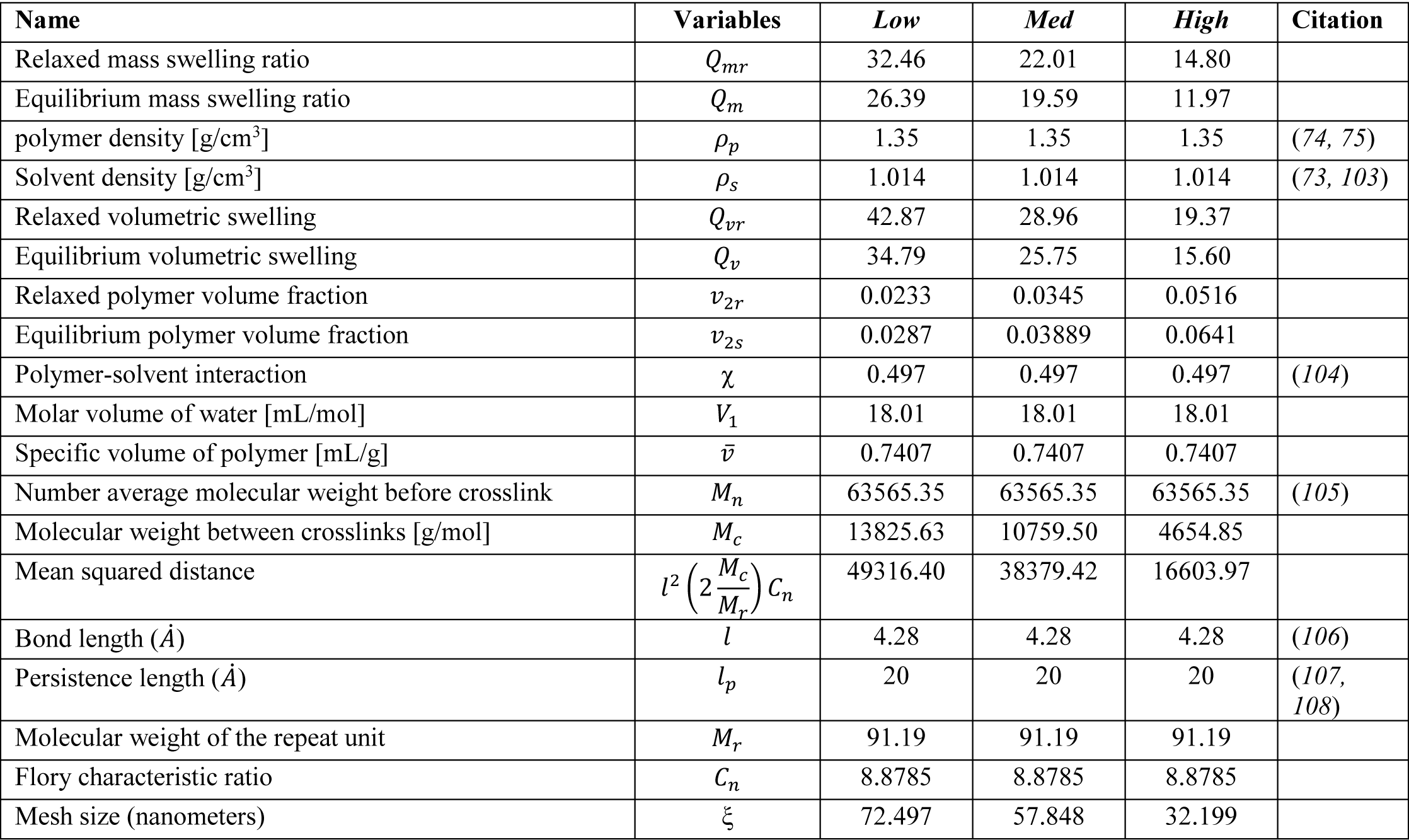
Parameters for hydrogel mesh calculations. Values with references were either taken directly or interpolated from literature.

