## supplemental only for "Mesenchymal stromal cell remodeling of a gelatin hydrogel microenvironment defines an artificial hematopoietic stem cell niche"

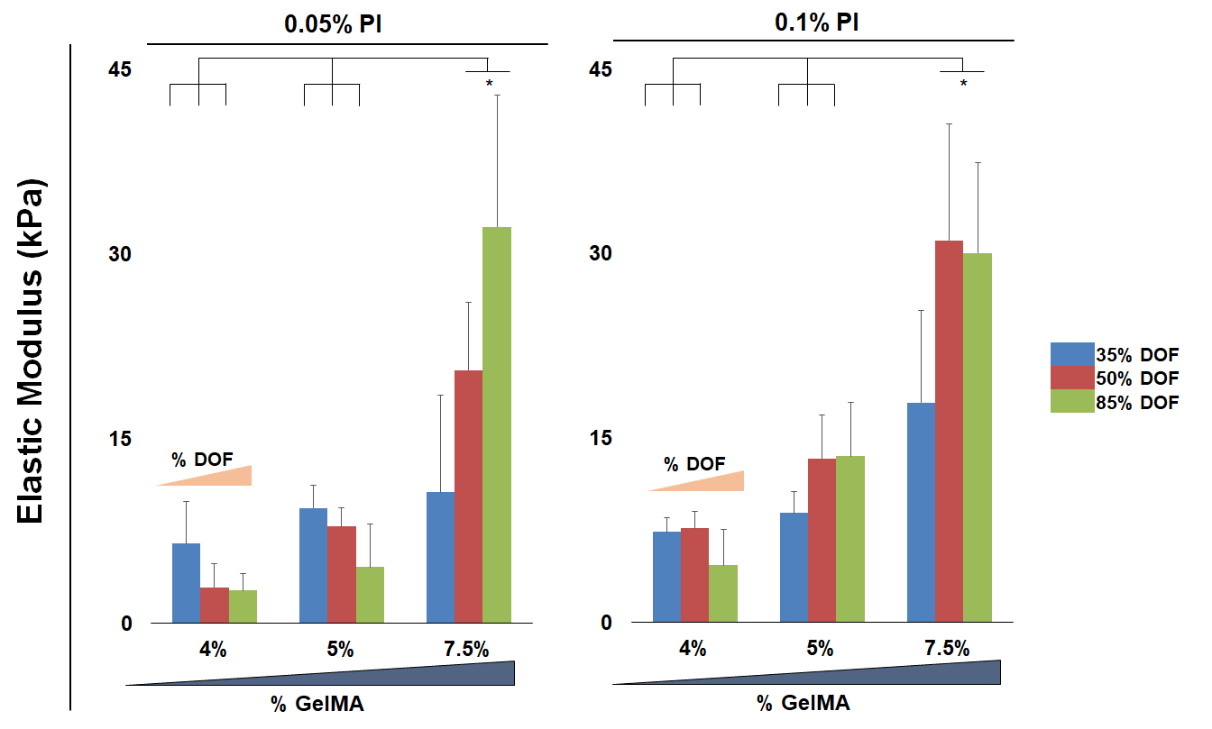

**Supplemental Figure 1.** Compressive mechanical analysis of the complete library of hydrogels. The degree of methacrylamide-functionalization was altered from 35, 50, 85%, and the content of GelMA was increased from 4, 5, 7.5%. * marks significantly different at the p<0.05 level. (**n=6**). **A**) Elastic modulus of hydrogels prepared with 0.05% photoinitiator. **B**) Elastic modulus of hydrogels prepared with 0.1% photoinitiator.

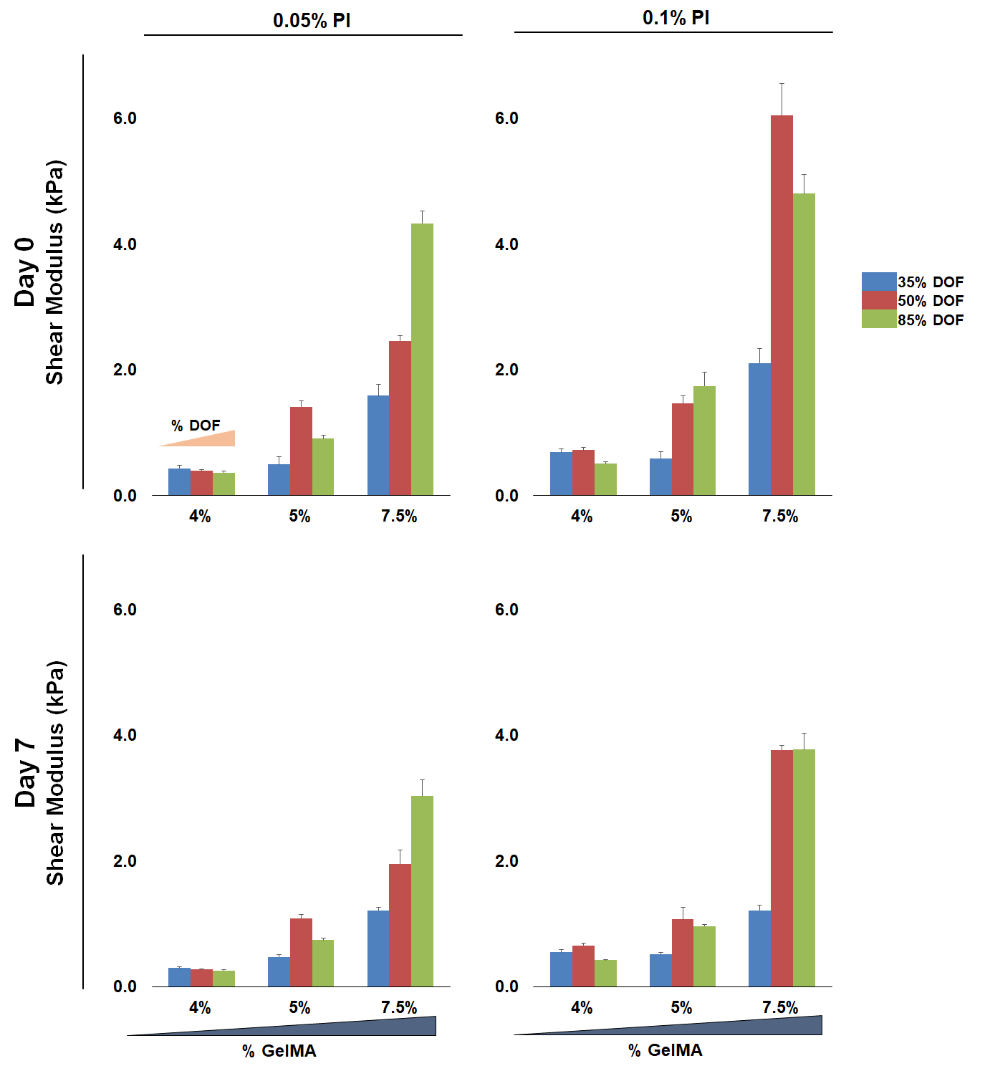

**Supplemental Figure 2.** Shear modulus characterization of the complete library of hydrogels. The degree of methacrylamide-functionalization was altered from 35, 50, 85%, and the content of GelMA was increased from 4, 5, 7.5%. (**n=6**). **A**) Shear modulus of hydrogels prepared with 0.05% photoinitiator, measured at Day 0 after hydration in PBS. **B**) Shear modulus of hydrogels prepared with 0.1% photoinitiator, measured at Day 0 after hydration in PBS. **C**) Shear modulus of hydrogels prepared with 0.05% photoinitiator, measured at Day 7. **D**) Shear modulus of hydrogels prepared with 0.1% photoinitiator, measured at Day 7.

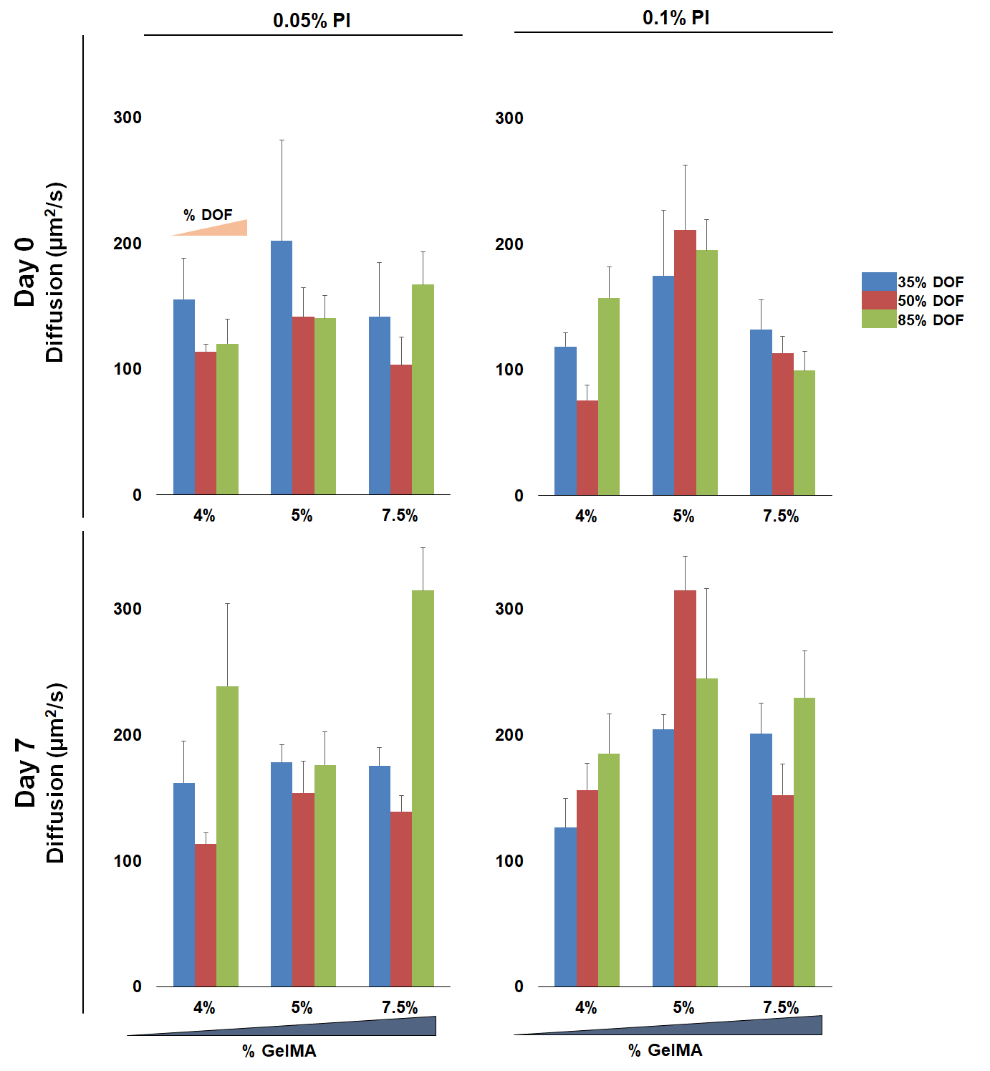

**Supplemental Figure 3.** The diffusion coefficient of water in each hydrogel was measured for the complete library of hydrogels. The degree of methacrylamide-functionalization was altered from 35, 50, 85%, and the content of GelMA was increased from 4, 5, 7.5%. (**n=6**). **A**) Diffusion coefficient of water in hydrogels prepared with 0.05% photoinitiator, measured at Day 0 after hydration in PBS. **B**) Diffusion coefficient of water in hydrogels prepared with 0.1% photoinitiator, measured at Day 0 after hydration in PBS. **C**) Diffusion coefficient of water in hydrogels prepared with 0.05% photoinitiator, measured at Day 7. **D**) Diffusion coefficient of water in hydrogels prepared with 0.1% photoinitiator, measured at Day 7.

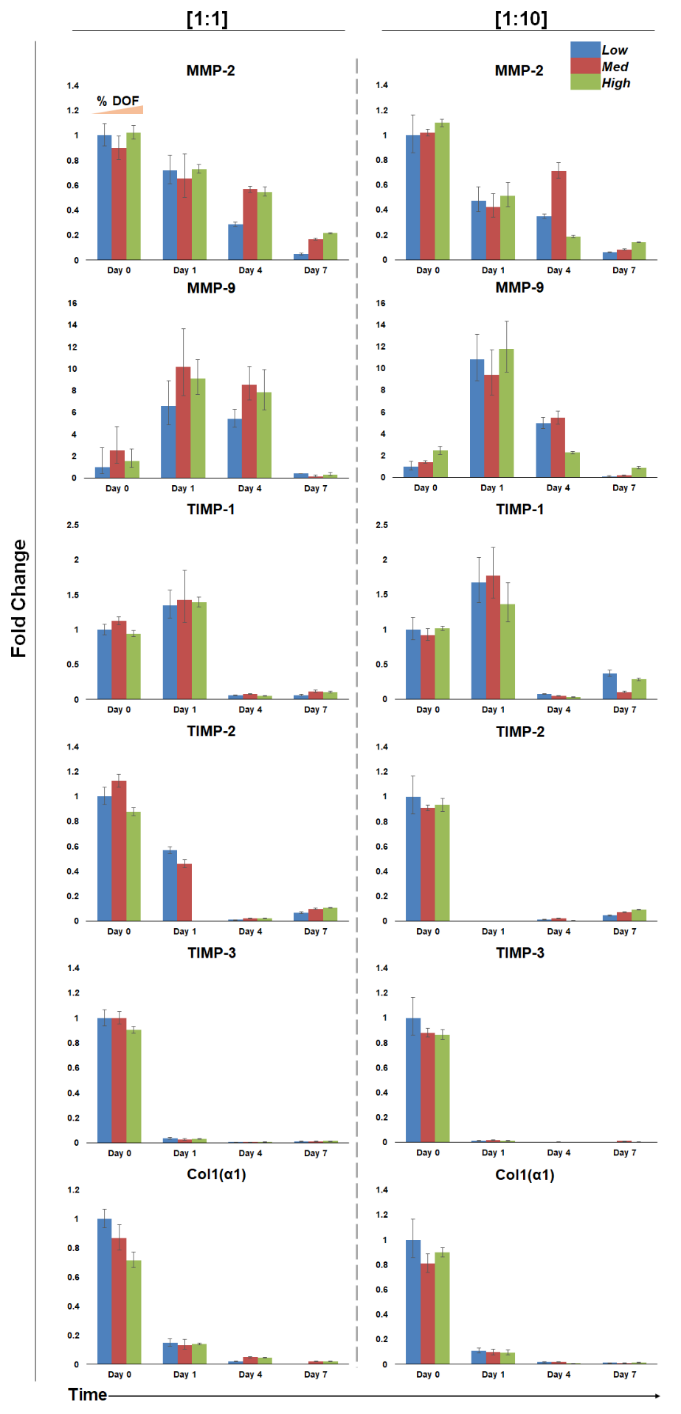

**Supplemental Figure 4.** Relative expression of matrix-associated genes in *Low*, *Med*, and *High* hydrogels seeded with 1x10^5^ (1:1) and 1x10^6^ MSCs/mL (1:10). GAPDH was the housekeeping gene, and expression was normalized to Day 0 of the low hydrogel condition. (**n=3-6**). **A**) Relative gene expression of hydrogels seeded with 1x10^5^ MSCs/mL (1:1) **B**) Relative gene expression of hydrogels seeded with 1x10^6^ MSCs/mL (1:10).

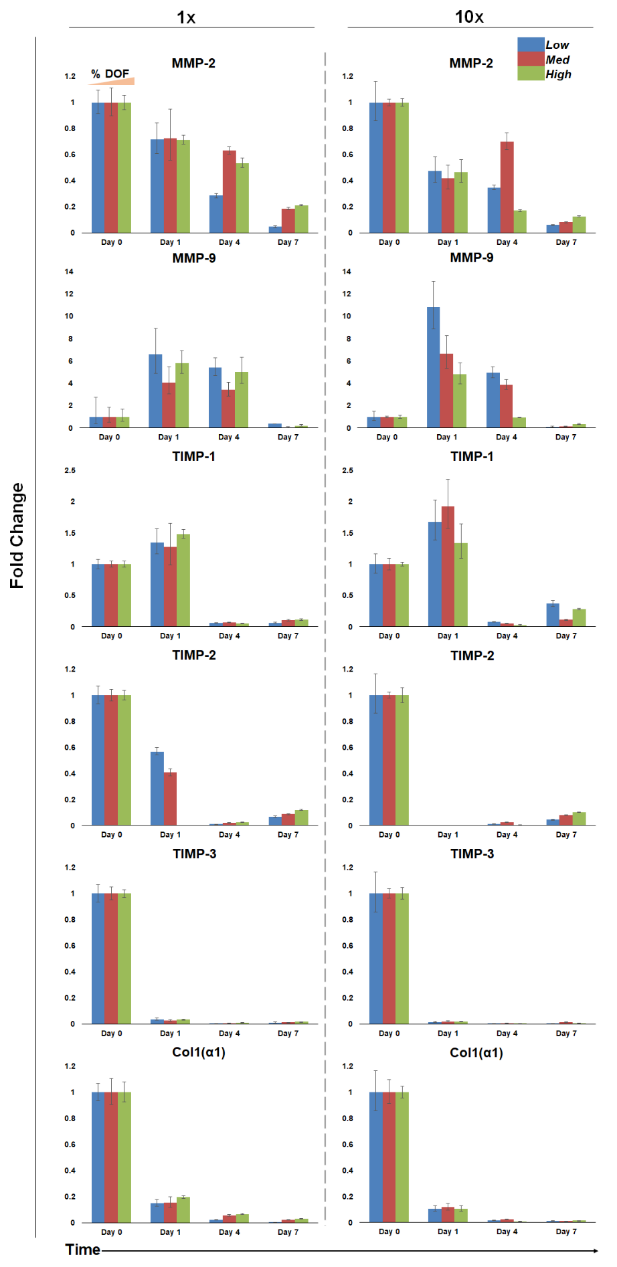

**Supplemental Figure 5.** Relative expression of matrix-associated genes in *Low*, *Med*, and *High* hydrogels seeded with 1x10^5^ (1:1) and 1x10^6^ MSCs/mL (1:10). GAPDH was the housekeeping gene, and expression was normalized to Day 0 of the specific hydrogel and seeding condition. (**n=3-6**). **A**) Relative gene expression of hydrogels seeded with 1x10^5^ MSCs/mL (1:1). **B**) Relative gene expression of hydrogels seeded with 1x10^6^ MSCs/mL (1:10).

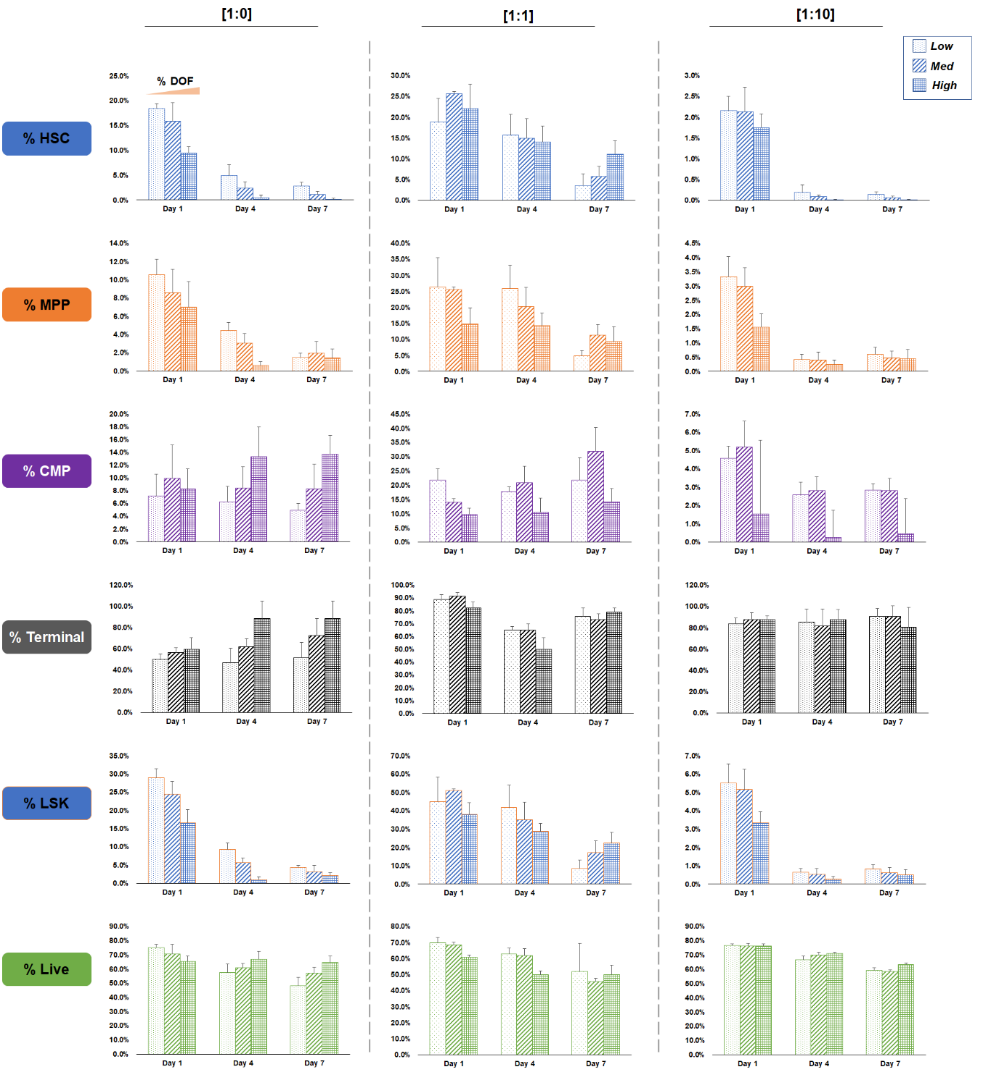

**Supplemental Figure 6.** Hematopoietic cell differentiation patterns within *Low*, *Med*, and *High* hydrogels with 0, 1x10^5^, and 1x10^6^ MSCs/mL (1:0, 1:1, and 1:10 respectively). Each cell type is expressed as a percentage of the overall hematopoietic cell population. (**n=6**). **A**) Hematopoietic stem cell population, comprised of long-term and short-term HSCs. **B**) Multipotent progenitor cells. **C**) Common myeloid progenitor cells. **D**) Terminal cell population, comprised of all non-LSK hematopoietic cells. **E**) LSK cells comprised of LT-HSC, ST-HSCs, and MPPs. **F**) The live population of cells expressed as a percentage of live cells over the total population of all cells.

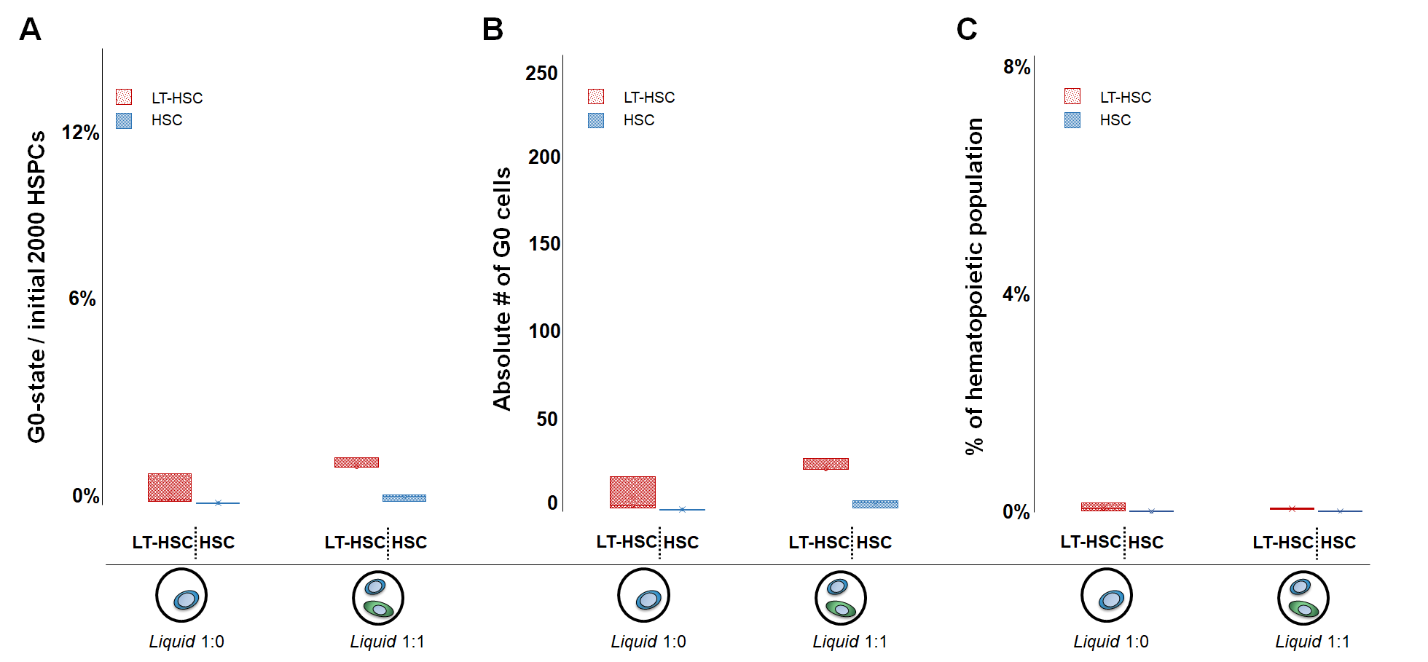

**Supplemental Figure 7.** Number and fraction of quiescent LT-HSCs and HSCs in liquid culture containing either 1x10^5^ HSPCs/mL alone (1:0), or a mixture of 1x10^5^ HSPCs/mL and 1x10^5^ MSCs/mL (1:1). (n=3). **A)** Number of quiescent LT-HSC or HSC, shown as a percentage of the initial 2000 HSPCs seeded. **B)** The absolute number of quiescent (G0) LT-HSC and HSCs within each sample (both populations are highest in the *High* variant at 1:1 seeding). **C)** Percentage of quiescent cells calculated vs. the total number of hematopoietic lineage positive cells after 7 days in culture (including hematopoietic progeny that arise during the culture period).

**Supplemental Table 1.**

Forward and reverse primer sequences for matrix associated genes. Sequences were made using NCBI Primer-blast or taken from literature.

| **Name** | **Accession #** | **Forward** | **Reverse** | **Citation** |
| --- | --- | --- | --- | --- |
| Col1(α1) | NM_007742.4 | CGACCTCAAGATGTGCCACT | GACGGCTGAGTAGGGAACAC | (*99*) |
| MMP-2 | NM_008610.3 | CCTGACCTGGACCCTGAAAC | TCCCAGCGTCCAAAGTTGAT | (*99*) |
| MMP-9 | NM_013599.4  PB#: 31560795c2 | GGACCCGAAGCGGACATTG | CGTCGTCGAAATGGGCATCT | (*100-102*) |
| TIMP-1 | NM_001044384.1 | TGGCATCTGGCATCCTCTTG | GGTCTCGTTGATTTCTGGGGA | (*99*) |
| TIMP-2 | NM_011594.3 | TATCTACACGGCCCCCTCTT | TCCCAGGGCACAATGAAGTC | (*99*) |
| TIMP-3 | NM_011595.2 | TCTCTGGACCGACATGCTCT | GCGTTGCTGATGCTCTTGTC | (*99*) |
| GAPDH | NM_001289726.1 | TGTGTCCGTCGTGGATCTGA | CCTGCTTCACCACCTTCTTGA | (*33*) |

**Supplemental Table 2.**

Parameters for hydrogel mesh calculations. Values with references were either taken directly or interpolated from literature.

| **Name** | **Variables** | ***Low*** | ***Med*** | ***High*** | **Citation** |
| --- | --- | --- | --- | --- | --- |
| Relaxed mass swelling ratio | $Q_{mr}$ | 32.46 | 22.01 | 14.80 |  |
| Equilibrium mass swelling ratio | $Q_{m}$ | 26.39 | 19.59 | 11.97 |  |
| polymer density [g/cm^3^] | $\rho_{p}$ | 1.35 | 1.35 | 1.35 | (*74, 75*) |
| Solvent density [g/cm^3^] | $\rho_{s}$ | 1.014 | 1.014 | 1.014 | (*73, 103*) |
| Relaxed volumetric swelling | $Q_{vr}$ | 42.87 | 28.96 | 19.37 |  |
| Equilibrium volumetric swelling | $Q_{v}$ | 34.79 | 25.75 | 15.60 |  |
| Relaxed polymer volume fraction | $v_{2r}$ | 0.0233 | 0.0345 | 0.0516 |  |
| Equilibrium polymer volume fraction | $v_{2s}$ | 0.0287 | 0.03889 | 0.0641 |  |
| Polymer-solvent interaction | χ | 0.497 | 0.497 | 0.497 | (*104*) |
| Molar volume of water [mL/mol] | $V_{1}$ | 18.01 | 18.01 | 18.01 |  |
| Specific volume of polymer [mL/g] | $\bar{v}$ | 0.7407 | 0.7407 | 0.7407 |  |
| Number average molecular weight before crosslink | $M_{n}$ | 63565.35 | 63565.35 | 63565.35 | (*105*) |
| Molecular weight between crosslinks [g/mol] | $M_{c}$ | 13825.63 | 10759.50 | 4654.85 |  |
| Mean squared distance | $l^{2}\left( 2\frac{M_{c}}{M_{r}} \right)C_{n}$ | 49316.40 | 38379.42 | 16603.97 |  |
| Bond length ($\dot{A}$) | $l$ | 4.28 | 4.28 | 4.28 | (*106*) |
| Persistence length ($\dot{A}$) | $l_{p}$ | 20 | 20 | 20 | (*107, 108*) |
| Molecular weight of the repeat unit | $M_{r}$ | 91.19 | 91.19 | 91.19 |  |
| Flory characteristic ratio | $C_{n}$ | 8.8785 | 8.8785 | 8.8785 |  |
| Mesh size (nanometers) | ξ | 72.497 | 57.848 | 32.199 |  |
